# Single-cell RNA-seq analysis of human iPSC-derived motor neurons resolves early and predictive ALS signatures

**DOI:** 10.1101/2020.04.27.064584

**Authors:** Ritchie Ho, Michael J. Workman, Pranav Mathkar, Kathryn Wu, Kevin J. Kim, Jacqueline G. O’Rourke, Mariko Kellogg, Valerie Montel, Maria G. Banuelos, Olubankole Aladesuyi, Sandra Diaz Garcia, Daniel Oheb, Steven Huang, Irena Khrebtukova, Lisa Watson, John Ravits, Kevin Taylor, Robert H. Baloh, Clive N. Svendsen

## Abstract

Induced pluripotent stem cell (iPSC) derived neural cultures from amyotrophic lateral sclerosis (ALS) patients can reflect disease phenotypes targetable by treatments. However, widely used differentiation protocols produce mixtures of progenitors, neurons, glia, and other cells at various developmental stages and rostrocaudal neural tube segments. Here we present a methodology using single-cell RNA sequencing analysis to distinguish cell type expression in C9orf72 ALS, sporadic ALS, control, and genome-edited cultures across multiple subjects, experiments, and commercial platforms. Combinations of HOX and developmental gene expression with global clustering classified rostrocaudal, progenitor, and mantle zone fates. This demonstrated that iPSC-differentiated cells recapitulate fetal hindbrain and spinal cord development and resolved early, reproducible, and motor neuron-specific signatures of familial and sporadic ALS. This includes downregulated ELAVL3 expression, which persists into disease endstages. Single-cell analysis thus yielded predictive ALS markers in other human and mouse models which were otherwise undiscovered through bulk omics assays.

## Introduction

The ability to analyze global gene expression in single cells within bulk tissues shifts the paradigm for understanding molecular mechanisms underlying cellular, tissue, and organismal physiology. Numerous recent studies have utilized single-cell RNA-seq (scRs) technology in primary human tissue as well as animal models (Keren-Shaul et al., 2017; Sala Frigerio et al., 2019; Stegle et al., 2015; Wagner et al., 2016) to survey and catalog cell identities (Macosko et al., 2015), developmental trajectories (Qiu et al., 2017), and physiological perturbations associated with disease (Mathys et al., 2019). scRs can also interrogate human induced pluripotent stem cell (iPSC)-based models of disease, such as in cases of familial and sporadic Parkinson’s disease (Lang et al., 2019).

Here, we applied scRs to understand the heterogeneous and cell type-specific nature of disrupted physiology in human iPSC-based models of amyotrophic lateral sclerosis (ALS), a neurodegenerative disorder characterized by cortical and spinal motor neuron (MN) death that results in weakness and paralysis of voluntary muscles (Ragagnin et al., 2019; Swinnen and Robberecht, 2014). While numerous molecular pathways and cell types associated with ALS have been described, definitive mechanisms responsible for MN degeneration remain elusive (Taylor et al., 2016). The vast majority of ALS cases are sporadic, with no known genetic link. In familial cases, ALS can be traced to a set of genetic mutations, for instance in the genes for SOD1 and C9orf72. However, symptom onset for both familial and sporadic ALS varies across body regions, thereby compounding difficulty in discerning disease etiology. Despite this variation, common clinical presentations are observed across familial and sporadic cases, suggesting that molecular features may converge across ALS patients. These features may encompass common mechanisms underlying MN degeneration that could be detected in patient iPSC models.

The ability to isolate robust disease signatures at single-cell resolution in iPSC disease models remains a challenging endeavor in many aspects. First, droplet based scRs, as a recently developed technology with lower sequencing depth compared to bulk RNA-seq, is vulnerable to technical confounders such as dropout, sensitivity to experimental batch effects, and variation across technological and commercial platforms (Hicks et al., 2018; Luecken and Theis, 2019). Second, iPSC-differentiated tissues do not recapitulate defined, mature, and adult-like states (Ho et al., 2016; Stein et al., 2014), which may reduce the fidelity of signals representing dysfunctional physiologies experienced by in vivo tissues in late onset diseases. Finally, iPSC-based modeling of nongenetic forms of disease requires a large number of subject lines to meaningfully represent populations of genetically diverse individuals; however, processing many samples within one experimental batch remains restricted by current costs and technological limitations of scRs.

We addressed the technical challenges of scRs of iPSC disease models by profiling multiple subject lines across multiple experimental batches and two scRs commercial platforms and performing meta-analysis to discern reproducible, cell type-specific, and early gene expression signatures of ALS. We combined supervised annotation of cell types by developmental gene markers with unsupervised global clustering to identify individual cells along the rostrocaudal axis of the mammalian body, as well as progenitor and postmitotic neural subtypes, which arise in complex cultures during in vitro differentiation. To isolate cell type-specific effects of the C9orf72 hexanucleotide repeat expansion (HRE) in ALS, we compare gene expression changes between C9orf72 HRE ALS cells to control subject cells as well as genome-edited isogenic cells. We also performed gene expression comparisons between sporadic ALS and control conditions and observed common changes occurring in C9orf72 ALS conditions. Highlighting the efficacy of this rigorous scRs approach, we uncover early signatures that persisted throughout early to end stages of disease in both familial and sporadic ALS cases as well as in animal models. Furthermore, the signatures discovered through scRs could be applied to bulk transcriptomic and proteomic data sets to accurately classify ALS from non-ALS cases. Our results suggest that these signatures are potentially causative of disease and highlight their value as predictive biomarkers that will aid future studies aiming to better characterize and treat ALS.

## Results

### Production of control, sporadic, C9orf72 ALS and isogenic iPSC derived MNs

MNs were differentiated from iPSC lines reprogrammed from either fibroblasts or peripheral blood monocytes from four healthy subjects: 0083, 0179, 0025, and 0465, two sporadic ALS subjects: 2XWC and 8BRM, and four familial ALS subjects with C9orf72 HRE: 0028, 0029, 0052, and 6ZLD (Table S1 and Figure 1A). To isolate C9orf72 HRE effects from inherent genetic variability, isogenic patient lines were established from two C9orf72 HRE lines (0029 and 0052) using CRISPR-Cas9-mediated gene editing to remove the HREs (Table S1 and Figure S1A). Successfully edited iPSC clones were kayotypically normal (Figure S1B-E), and, retained the ability to differentiate into MNs over a 30 day in vitro differentiation protocol (Yang et al., 2013) at a comparable efficiency to their parental C9orf72 HRE cell lines (Figure S2A). Removal of the repeat expansions resulted in two-fold increase in expression of all C9orf72 transcript variants back to levels observed in normal controls (Figures S2B and S2C) and eliminated the presence of RNA foci comprising sense and antisense sequences of the HRE (Figures S2D and S2E). Furthermore, the presence of polyGP dipeptide repeats, which accumulated in C9orf72 HRE ALS subject MN cultures over time (Figure S2F), were reduced to control subject levels (Figure S2G). The generation of isogenic MN cultures thus enabled the direct attribution of molecular phenotypes to the HRE in their parental C9orf72 ALS subject lines.

**Figure 1.**
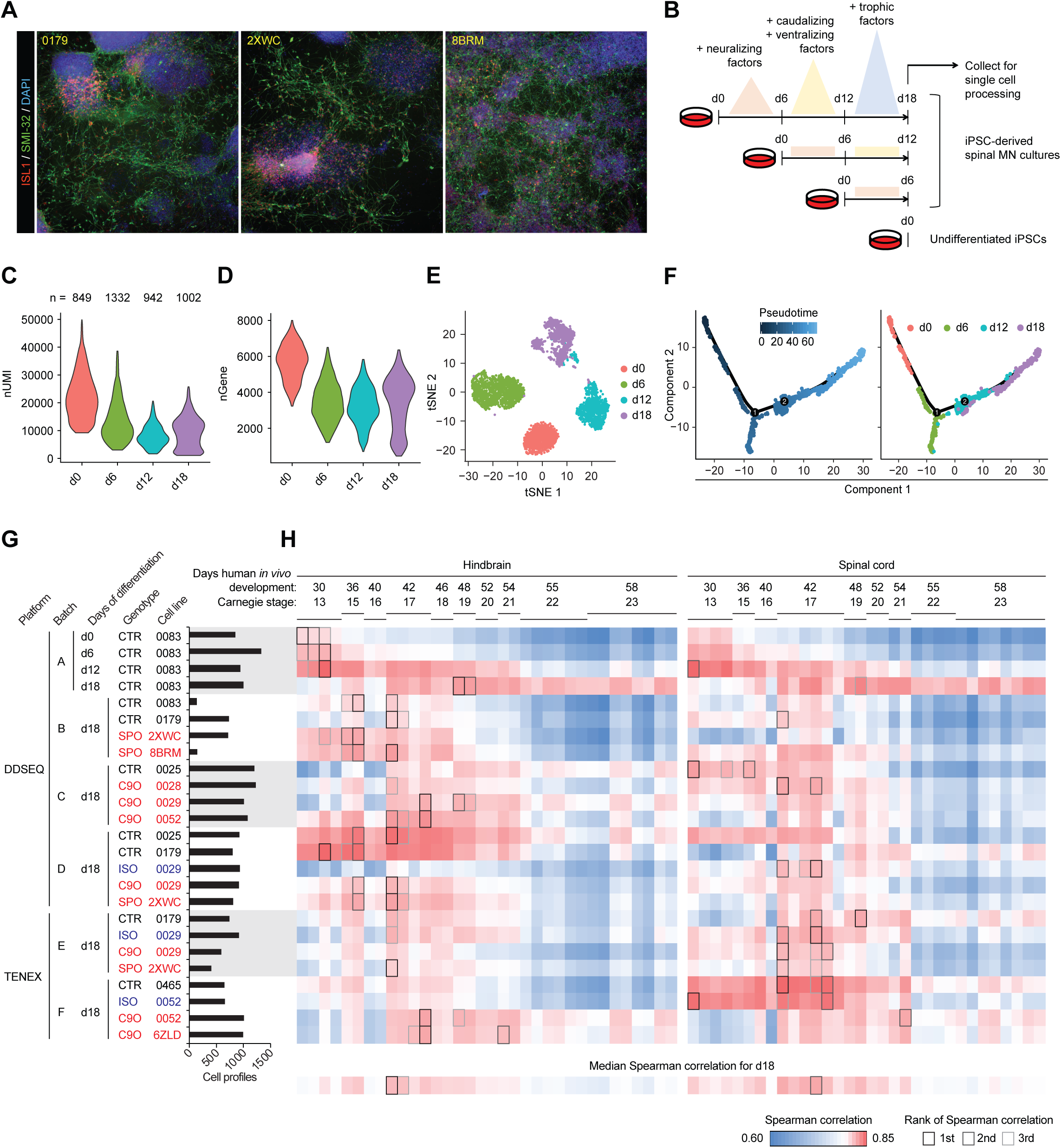
iPSC-MN cultures recapitulate developmental gene expression patterns. A. Immunostaining day 18 cultures from three individual subject lines for the expression of MN markers ISL1 and SMI-32. B. Experiment schematic for analyzing the time course of MN differentiation over the course of 18 days. C - D. Violin plots indicate refinement of gene expression programs during MN differentiation. C depicts the number of unique molecular identifiers (nUMI), and D depicts the number of detectable genes per time point. E. tSNE of spinal MN differentiation time course samples. F. Monocle analysis projects samples into a pseudo-time axis consistent with the order of time points. G. Histogram displays meta data for all samples profiled in this study. This data is also presented in Figure S4. H. UMI counts were summed for each gene across the expression matrix for each sample, thereby simulating bulk RNA-seq data. Simulated bulk gene expression profiles for each sample were correlated to bulk RNA-seq gene expression profiles from fetal hindbrain and spinal cord at various Carnegie stages analyzed in de Kovel et al., 2017. The median Spearman correlation was calculated for each column of day 18 samples and displayed along the bottom row. For each row of pairwise correlations, the top three ranked correlations are outlined, indicating which sample from de Kovel et al., 2017 most globally similar to each sample analyzed by single-cell RNA-seq.

### iPSC-derived MN cultures recapitulate developmental gene expression patterns

Next, the differentiation of iPSC-MNs were characterized at the single-cell level using the Illumina® Bio-Rad® SureCell™ WTA 3’ Library Prep Kit for the ddSEQ™ System for one control line (0083) using a more rapid 18 day differentiation protocol that produces cranial and spinal MNs and interneurons (Maury et al., 2015: Figure 1B). Consistent with previous observations (Efroni et al., 2008), pluripotent cells undergo a reduction in overall transcriptional activity upon differentiation suggesting a refinement of transcriptional programs from the pluripotent to progenitor state (Figures 1C and 1D). Interestingly, the median of unique molecular identifiers (nUMIs) per cell increased between days 12 and 18 reflecting perhaps a state of specialized physiology and functions. Global clustering resolved each time point into distinct clusters, where day 12 and day 18 populations further resolved into subpopulations (Figure 1E). Pseudotime analysis through Monocle (Qiu et al., 2017) of cells from all time points arranged each time course in the expected order of progressively differentiating cell states (Figure 1F). 20 marker genes for spinal MN development and maturation (Ho et al., 2016) were expressed along the pseudotime axis in a pattern consistent with fetal-like tissues derived in vitro from iPSCs (Figure S3A).

### iPSC-MN cultures globally resemble fetal hindbrain and spinal cord

We next performed scRs on MN cultures from several ALS and control subject lines at 18 days of differentiation in order to establish a pool of single cells we could use to determine regional specificity as well as the presence of ALS signatures (Table S1 and Figure 1G). Because only a finite amount of samples could be captured and processed within each experiment, we collected samples across six batches of differentiation (A-F). We also aimed to establish the robustness of any signal across two different scRs platforms: the Illumina Bio-Rad Single-Cell Sequencing Solution (DDSEQ) and the 10X (TENEX) Genomics Chromium (Table S1 and Figure 1G). Immunostaining and quantification of day 18 cultures indicated no significant differences in ISL1 and SMI-32 positive MNs between ALS and control suggesting that an overt disease phenotype such as cell death has not manifested at this relatively early differentiation time point (Figure S3B) as shown in previous studies (Fujimori et al., 2018; Sareen et al., 2013). In total, we analyzed 21,702 cells that passed quality control filters. To gauge the developmental and maturation states of these cultures, we correlated their expression profiles to previous data sets characterizing spinal MN maturation gene expression (Ho et al., 2016) (Figures S4A and S4B) and to a bulk RNA-seq dataset of human fetal hindbrain and spinal cord tissue ranging from Carnegie stages 13 to 23 (de Kovel et al., 2017) (Figure 1H). By 18 days, iPSC-derived cells showed transcriptional states that most globally resemble fetal hindbrain and spinal cord tissue at Carnegie stage 17, or about 42 days of in vivo development.

In order to establish the rostrocaudal identity of individual cells, we next focused on the family of homeobox transcriptional regulators of morphological patterning, the HOX genes. Based on previous genetic studies (Di Bonito et al., 2013; Lippmann et al., 2015; Philippidou and Dasen, 2013), we composed a model for relative HOX gene expression along the rostrocaudal axis ranging from rhombomeres two to eight of the developing hindbrain and cervical to caudal segments of the spinal cord (Figures 2A, 2B, and Table S2A). We observed that RNA expression levels of each HOX gene in the fetal hindbrain and spinal cord tissue samples from de Kovel et al., 2017 are consistent with our model, and classification of segment identity based on the highest correlation for each sample resolves the sample types (Figures 2B and 2C). While correlation of bulk profiles from day 18 iPSC-MN cultures suggested that the cultures globally resembled hindbrain more than spinal cord, we hypothesized that some rare cells may have differentiated into more caudal identities. We therefore applied this classification approach for each single cell in the day 18 cultures. These results indicated that while a majority of cells (33.19%) were not assigned (NA) categories at the single-cell level, either due to a lack of any detectable HOX gene expression or failure to meet the correlation cutoff, the second majority of cells (25.75%) were classified as rhombomere eight, and a third majority of cells (10.78%) were classified as the cervical segment (Figure 2C and Table S2C). Notably, there were some cells classified as brachial (1.32%) and thoracic (1.40%) segments, suggesting that the 18 day protocol can achieve differentiation into cell types within spinal cord that reflect the upper limb sites of disease onset for most subjects represented in this study (Table S1). However, there were no cells classified as lumber segment, possibly due to the early differentiation time point of these day 18 cultures.

**Figure 2.**
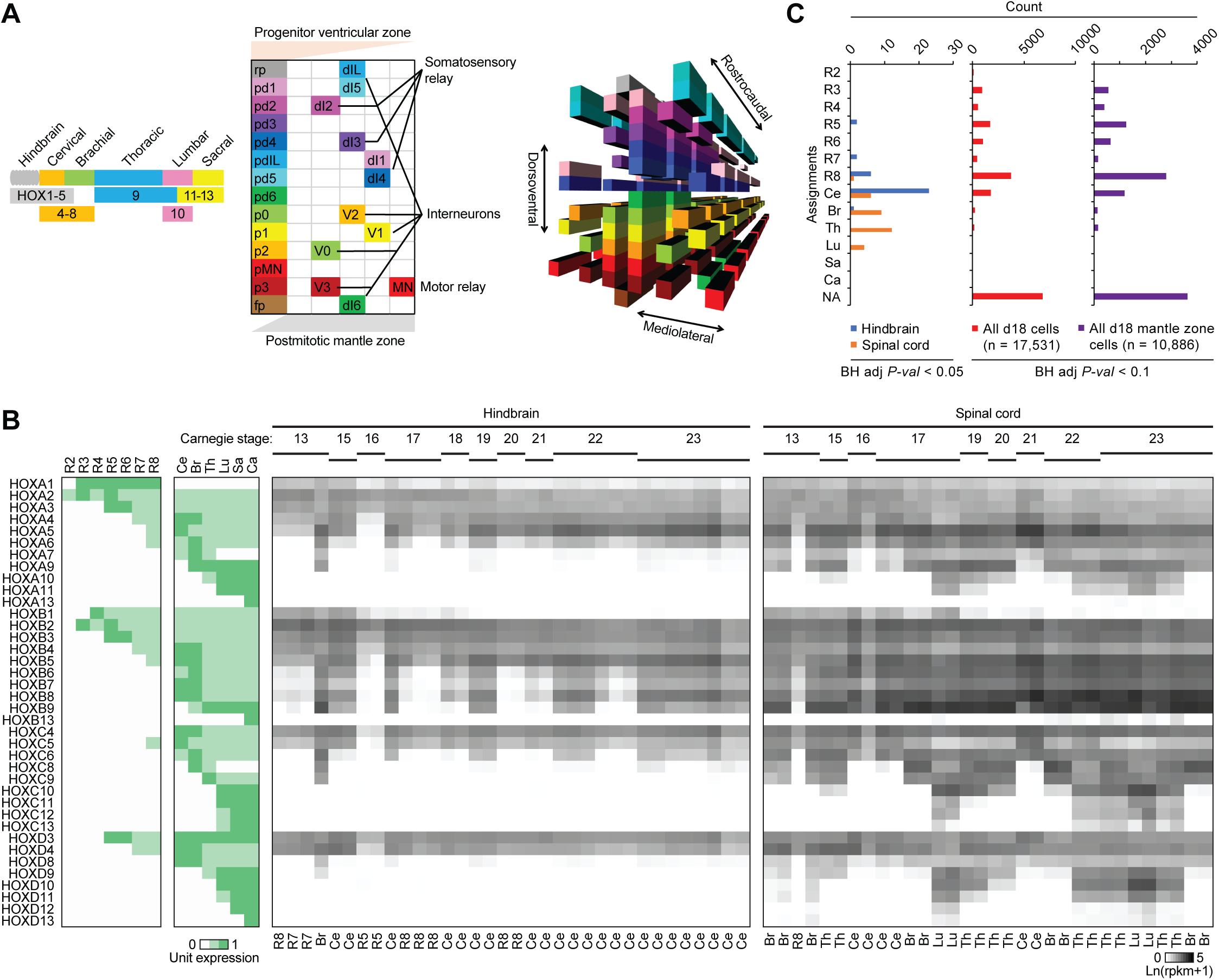
iPSC-MN cultures globally resemble rhombomere eight and cervical spinal cord. A. Models of cell type organization along representative hindbrain and spinal cord segments. Left, HOX gene expression patterns determine rostrocaudal axis identities, adapted and simplified from Di Bonito et al., 2013; Lippmann et al., 2015; and Philippidou and Dasen, 2013. Center, dorsoventral progenitors in the ventricular zone (VZ) and postmitotic lineages localized to distinct regions in the gray matter mantle zone (MZ) of the spinal cord, adapted and simplified from Alaynick et al., 2011 and Lu et al., 2015. Right, registration of all cell types present within the hindbrain and spinal cord based on HOX and lineage-specific transcription factor expression. B. Left: Unit expression matrix of HOX genes that developmentally determine hindbrain and spinal cord segment identity. R2-8: rhombomeres 2-8, Ce: cervical, Br: brachial, Th: thoracic, Lu: lumbar, Sa: sacral, Ca: caudal. Right: Expression heatmap of HOX genes in fetal hindbrain and spinal cord at various Carnegie stages analyzed in de Kovel et al., 2017. Labels below each sample column indicate the hindbrain or spinal cord segment that each sample most resembles, based on the highest Spearman correlation to the expression patterns for each segment shown in the unit expression matrix and with Benjamini-Hochberg adjusted *P*-values < 0.05. C. Histogram of segment identities assigned to fetal hindbrain samples, fetal spinal cord samples, and individual cells analyzed from day 18 cultures based on the highest Spearman correlation to the expression patterns for each segment shown in the unit expression matrix and with Benjamini-Hochberg adjusted *P*-values meeting the indicated thresholds. NA: not assigned. See also Figure S5, and Table S2.

### Developmental gene expression profiles and global clustering classify ventricular zone (VZ) progenitor and mantle zone (MZ) postmitotic neuronal identities

The induction of neural differentiation occurs after embryonic regionalization of the anterioposterior axis (Metzis et al., 2018). It involves the specific expression of 105 genes encoded in a two-dimensional coordinate system of morphogen gradients that regulate the dorsoventral and mediolateral axes and the progression of neural progenitors to postmitotic neurons in a representative spinal cord segment (Alaynick et al., 2011; Lu et al., 2015) (Figures 2A, S5A, and Table S2B). We next sought to resolve individual neural identities using these genes in day 18 cultures. We focused on correlating each cell type in our model with one another based on these 105 genes and demonstrated that these profiles can systematically distinguish each identity (Figure S5B). Assignment of individual cells along the 18 day differentiation to either VZ progenitors or MZ postmitotic neurons illustrated a cell fate progression consistent with the functions of the morphogenic components used during induction (Figures 1B, 3A, and Table S2C). Notably, very few astrocytes were seen (Figure 3A) indicating that the rapid 18 day differentiation did not enact a glial program.

**Figure 3.**
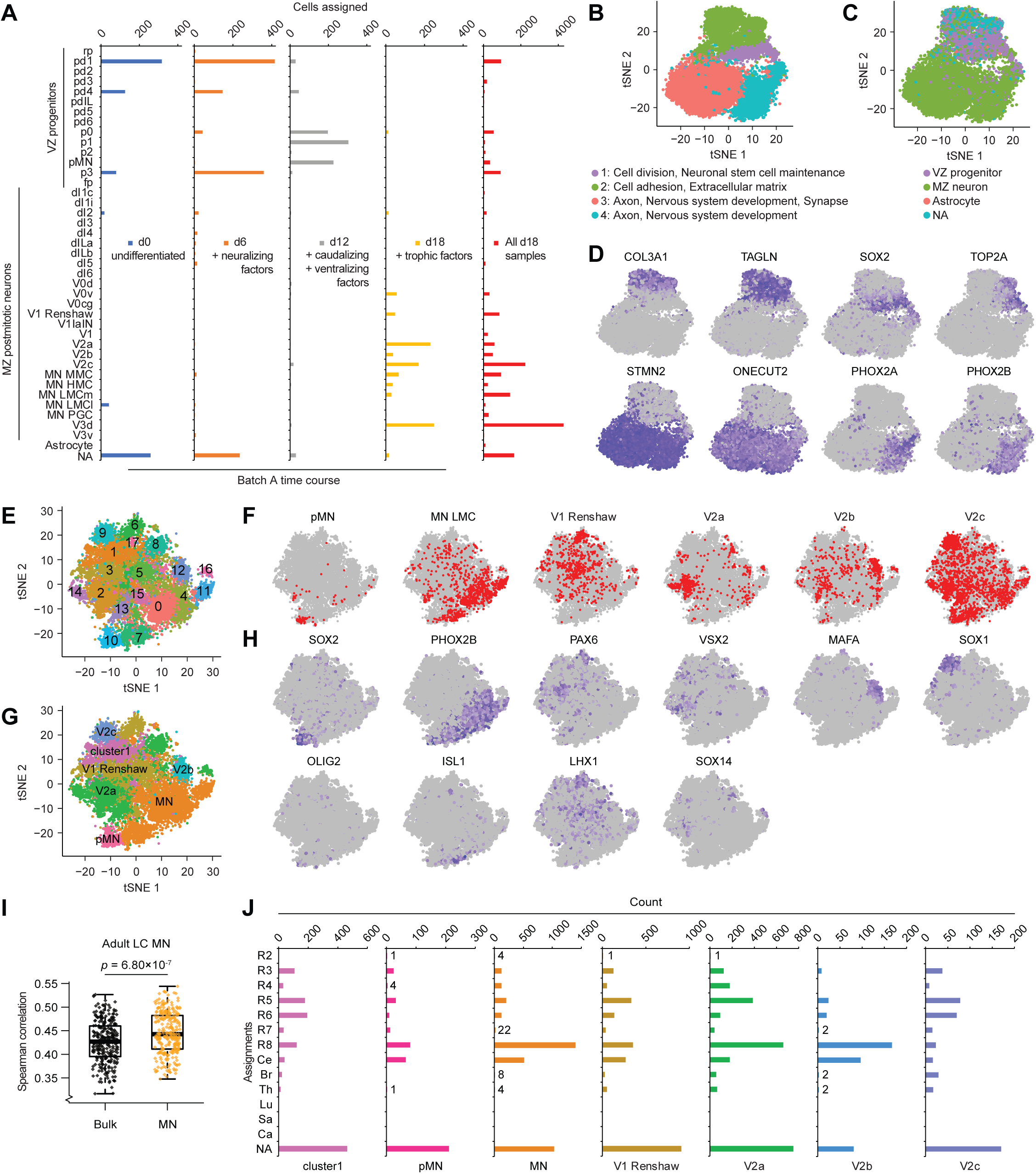
Developmental gene expression profiles and global clustering classify VZ progenitor and MZ postmitotic neuronal identities. A. Based on the expression profile of 105 developmental genes (Table S2B), individual cells from the time course of differentiation over 18 days were classified as identities belonging to the VZ, MZ, astrocyte, or not assigned (NA). B. tSNE showing global clustering of 17,531 cells from day 18 cultures across experimental batches, which by Seurat determined four clusters indicated by four colors. Significantly differentially expressed genes in each cluster relative to the other clusters were analyzed for enriched GO terms, and these are displayed below the tSNE plot. C. The same tSNE as in B with each cell colored by classification based on 105 developmental genes. D. The same tSNE as in B with each cell colored by relative expression level of each indicated gene, which were determined to be among the most significantly differentially expressed genes in clusters 1 (SOX2, TOP2A), 2 (COL3A1, TAGLN), 3 (STMN2, ONECUT2), and 4 (PHOX2A, PHOX2B). E. tSNE showing global clustering of 10,866 cells classified into clusters 3 and 4 in B. Seurat analysis determined 18 clusters. F. The same tSNE as in E with each cell colored red if they were classified as the indicated cell type based on 105 developmental genes. G. The same tSNE as in E with each cell colored by their reclassification into clusters formed by merging the clusters identified in E based on enrichment for identities shown in F. This data is also presented in Figure 4. H. The same tSNE as in E with each cell colored by relative expression level of each indicated gene, which are regarded as distinct markers of each identity listed above each plot in F. I. Boxplots of Spearman correlation values between gene expression profiles of LC MNs from Ho et al., 2016; summed gene expression profiles across all cells for each day 18 sample (Bulk); and summed gene expression profiles across cells classified as MN in G. *P*-values were calculated from a paired, two-tailed t-test. n.s. = not significant, *p* > 0.05. J. Bar graph of segment identities assigned to individual cells within each of the seven clusters based on the highest Spearman correlation to the HOX expression patterns for each segment shown in the unit expression matrix (Table S2A) and with Benjamini-Hochberg adjusted *P*-value < 0.1. R2-8: rhombomeres 2-8, Ce: cervical, Br: brachial, Th: thoracic, Lu: lumbar, Sa: sacral, Ca: caudal, NA: not assigned. See also Figures S4, S5, and S6, and Table S2.

Additionally, we used unsupervised global gene expression profiles to unbiasedly cluster distinct identities present in day 18 cultures. However, dimensional reduction and projection using principal component analysis (PCA) and t-distributed stochastic neighbor embedding (tSNE) of raw expression data primarily separated cells based on single-cell technology platform (Figure S5C). Experimental batch effects were also evident for samples processed within the same platform. These preliminary data highlighted the need to normalize the scRs expression data prior to discovering common variations between ALS and control conditions. To this end, multi-canonical correlation analysis (MultiCCA) in Seurat (Butler et al., 2018) corrected for experimental batch and platform effects (Figure S5D). By optimizing clustering parameters to yield a maximum modularity value of all communities, this analysis revealed four major populations of cells that enriched for expression of genes associated with a variety of gene ontology (GO) terms (Figures 3B-D, S5E, and Table S2C). Altogether, these analyses enable the resolution of major populations present in this rapid differentiation protocol, revealing not only postmitotic neurons generated from iPSCs, but also persistent progenitors and another population of non-neuronal cells.

In order to specifically detect neuronal subtype signatures in these cultures, we repeated subpopulation detection by removing non-neuronal cells and progenitors and then performing a new batch correction and global cluster analysis. This analysis assigned 18 major populations of cells (Figures 3E, S5F, and Table S2C). Overlaying each of the VZ and MZ identities with the globally defined clusters revealed that six of the classification groups reconciled with the global clustering patterns. We therefore renamed them based on these observations (Figure 3F-3H and Table S2C). For example, cells assigned as MNs of the lateral motor column (MN LMC) were enriched in clusters 0, 4, 7, 11, 15, and 17, and these cells expressed the MN markers PHOX2B (Pla et al., 2008) and ISL1 (Liang et al., 2011). We therefore renamed this group MN hereafter (Figure 3G). Immunostaining cultures confirmed protein co-expression of PHOX2B with ISL1, and distinct expression of the V2a and V2c interneuron markers VSX2 and SOX1, respectively (Figure S6A). Overall, based on overlapping classifications and expression of key marker genes, subsets of the 18 populations were merged to produce seven major populations (Figure 3G and Table S2C).

We then assessed whether the cells classified as MN, when segregated from the rest of the day 18 culture cells, showed more of an adult MN expression profile than if all cells were analyzed in bulk. By correlating only the pooled MN expression profiles to our previously characterized data set (Ho et al., 2016), MNs were significantly more correlated to in vivo adult MNs (Figures 3I, S4C-E). By subsetting cells into seven populations, reanalysis of rostrocaudal identity based on HOX gene expression demonstrated that the distributions of hindbrain and spinal cord segments are largely consistent across all populations (Figure 3J). Notably, cluster 1, V1 Renshaw, V2a, and V2c populations contained a modest number of cells resembling brachial and thoracic identities. These results highlight the value of scRs in resolving cell types to enable more accurate measures of similarity between in vitro iPSC-derived models and in vivo adult cell types.

### Pooling sparse transcriptional changes detected by scRs defines cell type-specific ALS responses

Having defined these seven populations, we performed differential gene expression between ALS and control conditions. After dividing each population into ALS and control groups, a comparable number of cells remained for each condition (Figure 4A), supporting the results determined by protein immunostaining for MN markers at this time point (Figure S3B). Tracking the scRs platforms also demonstrated equal representation of ALS and control groups assayed within each platform (Figure 4A). There were sufficient numbers of MNs, V1 Renshaw, and V2a interneurons from each experimental batch to perform differential gene expression analysis. Conducting comparisons between ALS and control conditions (which included isogenic C9orf72 HRE-corrected lines) yielded genes called significantly differentially expressed (data not shown). However, latent categorical variables such as experimental batch and scRs technology platform effects mainly drove these differences, illustrating the pitfalls of performing differential gene expression analysis without accounting for these properties. Thus, we next applied a meta-analysis approach by conducting comparisons between ALS and control or isogenic samples within each experimental batch and catalogued genes called significant (Figure 4B and Tables S3-7). For each ALS to control comparison (sporadic ALS samples presented in orange, C9orf72 ALS samples in magenta, control samples in black, and isogenically corrected HRE samples in green), the list of significantly upregulated genes (enumerated in red) were intersected with all other ALS to control comparisons, and the red heatmap indicates the Jaccard index, a measure of overlap between gene sets (Figure 4B). A similar analysis was performed on the downregulated genes (enumerated in blue) and presented in the blue heatmap. Interestingly, the number and concordance of genes called significantly dysregulated were highly variable across several comparisons, including repeated comparisons performed between two subject lines across different experimental batches (Figure 4B and Table S6). This indicated that despite assaying the same genetic comparisons, batch effects are observed, which may have arisen either by distinct biological responses to repeated differentiation experiments or by distinct technical effects across sample processing, both within and across commercial scRs platforms. Furthermore, there was low concordance of dysregulated genes when C9orf72 HRE lines were compared directly to their isogenically corrected lines. This observation highlights a challenge in detecting a reproducible gene expression signature of the C9orf72 HRE using scRs analysis of iPSC models, even when genetic variation is controlled.

**Figure 4.**
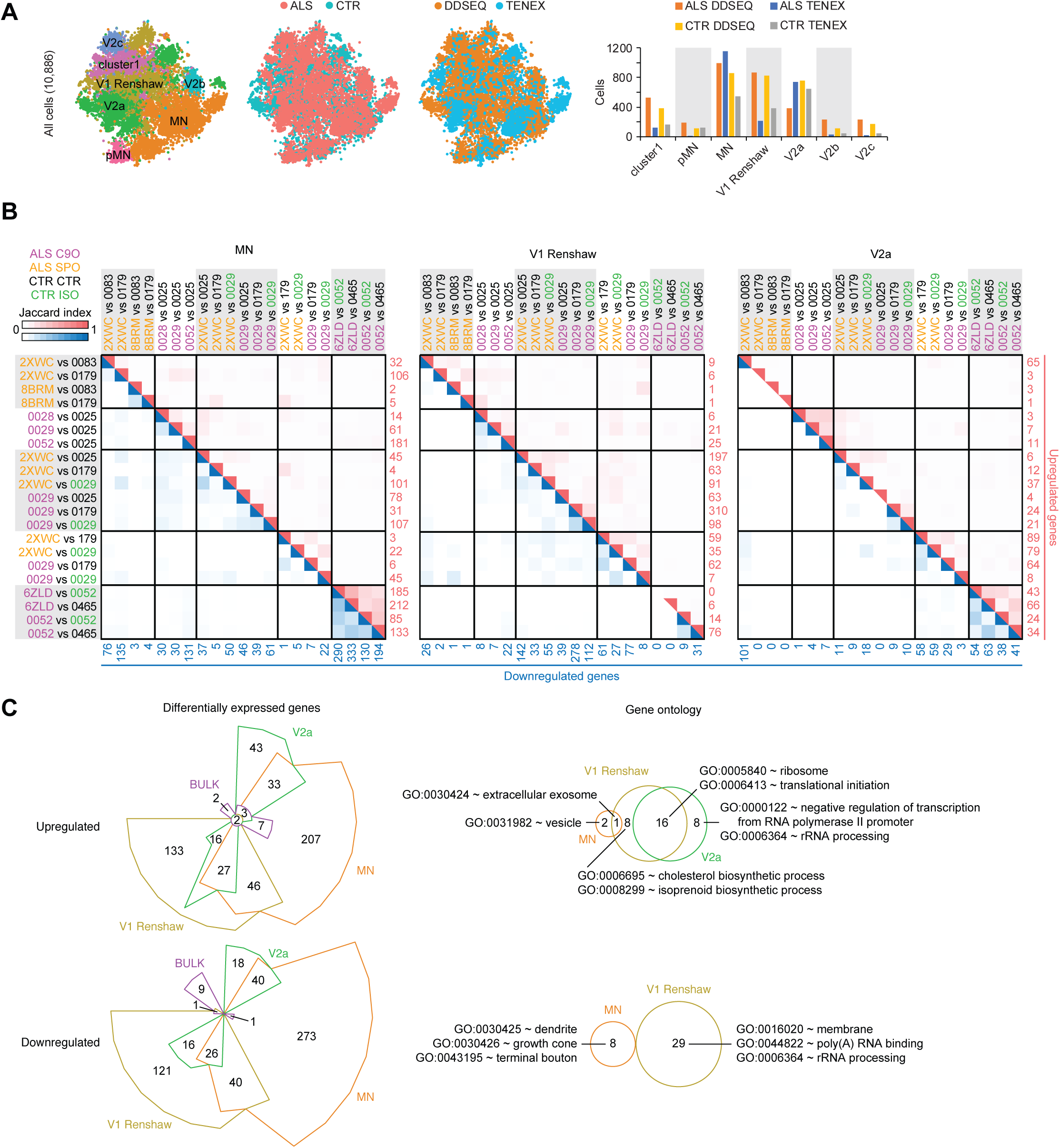
ALS enacts distinct gene expression changes in neuronal subtypes. A. Left to right: the same tSNE as in Figure 3G, with each cell colored by ALS or control (CTR) condition, colored by experimental platform, and bar graph quantifying the number of cells assigned to each class. B. Jaccard indices of overlapping gene sets across ALS to CTR comparisons within the neuronal subtypes MN, V1 Renshaw, and V2a. Each pairwise ALS to CTR comparison is shown along left and top axes, and the heatmap indicates the Jaccard index for each intersection of gene sets. The number of upregulated genes in each ALS to CTR comparison is printed in red along the right axis, and the number of downregulated genes in each ALS to CTR comparison is printed in blue along the bottom axis. Upregulated gene sets were only compared to other upregulated gene sets, and downregulated gene sets were only compared to other downregulated gene sets. The red and blue heat maps represent the Jaccard indices for intersections of upregulated and downregulated gene sets, respectively. C. Left: Differentially expressed genes between ALS and control conditions for neuronal subtypes MN, V1 Renshaw, V2a, and simulated bulk RNA-seq. Genes were categorized as upregulated or downregulated by ALS, and Chow-Ruskey diagrams present the intersection among each neuronal subtype as well as with bulk comparisons. Right: Euler diagrams present the intersection of GO terms enriched among genes that are reproducibly dysregulated by ALS uniquely within each neuronal subtype in the upregulated or downregulated categories. Representative GO terms are displayed from each overlapping or unique set. See Table S7H-M for full set of uniquely enriched GO terms. No GO terms were significantly enriched among genes upregulated or downregulated by ALS in bulk comparisons, and no GO terms were enriched among genes uniquely downregulated by ALS in V2a interneurons. See Tables S3-7 and Methods for details on how differentially expressed gene and GO lists were generated. See also Figure S6 and Tables S3, S4, S5, S6, and S7.

Given the sparseness of genes that were reproducibly discovered to be dysregulated across experimental batches, we next catalogued and pooled upregulated and downregulated genes called significant in at least two ALS to control or isogenic sample comparisons. This was done for the C9orf72 HRE ALS lines (12 comparisons) and the sporadic ALS lines (9 comparisons) (Tables S3, S4, and S5). Since our goal was to find early, convergent signatures across familial and sporadic forms of ALS, we respectively compared the extent of overlap between the upregulated and downregulated gene sets for each category between C9orf72 HRE and sporadic ALS conditions. Through hypergeometric testing, all comparisons indicated that the gene sets catalogued for both ALS conditions overlapped significantly (Figure S6B). We therefore combined the sparse set of differentially expressed genes from the C9orf72 HRE lines together with the sporadic ALS lines to amass gene sets large enough to pursue subsequent enrichment analyses. To this end, we catalogued and pooled genes called significant in at least two of the 21 ALS to control or isogenic sample comparisons drawn across all scRs experiments. With this approach, we generated a list of upregulated and downregulated genes for each of these three majority populations in our cultures (Tables S6 and S7A), and we compared these gene lists across all three populations (Figure 4C). Furthermore, we compared these gene lists to differentially expressed genes calculated by bulk analysis of all cells (Table S7A). This comparison demonstrated ALS can induce some overlapping but mostly distinct gene expression changes in each of the three iPSC-derived neuronal populations. Resolving cells into subpopulations was necessary to detect reproducibly disrupted genes, because analysis on the bulk expression profiles of the entire culture did not yield a high number of genes in either the upregulated or downregulated categories (Figure 4C and Table S7A).

GO analysis on the entire list of upregulated or downregulated genes from each cell type determined overlapping and distinct GO terms enriched among each list (Figure S6C and Table S7B-G). Analysis on the upregulated and downregulated genes that were unique to each cell type further refined GO terms (Figure 4C and Table S7H-M). Notably, components involved with translation and ribosomal subunits were commonly enriched among upregulated genes in all three neuronal cell types, but functions in cholesterol and isoprenoid synthesis were enriched among genes uniquely upregulated in V1 Renshaw interneurons. While translational components were also enriched among genes downregulated in all three neuronal cell types, components of neuronal processes including dendrite and growth cone were enriched among genes uniquely downregulated in MNs.

### ALS iPSC-MN cultures exhibit transcriptional changes detectable in postmortem ALS spinal MNs

We next tested the pathological relevance of these iPSC-MN defined gene sets by examining postmortem, adult spinal MNs. In previous work, we defined 52 co-expression modules using Weighted Gene Co-expression Network Analysis (WGCNA) (Zhang and Horvath, 2005) from laser capture micro-dissected MNs (LC MNs) from postmortem sporadic ALS and control subjects (Ho et al., 2016; Rabin et al., 2010), herein referred to as data set A. Some of these modules significantly correlated or anti-correlated to a principal component that distinguished sporadic ALS from control conditions. We systematically tested whether each list of upregulated or downregulated genes from MNs, V1 Renshaw, and V2a interneurons were enriched in each of the 52 previously defined WGCNA modules (Figure 5A). Markedly, the genes upregulated and downregulated by ALS in MNs were significantly enriched among modules that were respectively upregulated and downregulated by sporadic ALS in postmortem spinal MNs. This concordant response to ALS was not observed for V1 Renshaw and V2a interneurons. A repeated analysis between our scRs data set and an independent but similar postmortem study (Krach et al., 2018), herein referred to as data set B, demonstrated reproducibly concordant gene expression changes (Figure 5B). The robustness of networks defined in each of the postmortem data sets were also examined using module preservation z-statistics (Langfelder et al., 2011), which indicates the likelihood that the network structures of each module occurred by random chance. The most significantly overlapping modules, namely the Magenta, Midnightblue, Blue (Figure 5A), and Darkgreen modules (Figure 5B) possess the some of the most preserved network structures across data sets A and B (Figures S6D and S6E), suggesting they support critical functions in MNs. Importantly, dysregulation of these network genes in iPSC-MNs suggests that their disruption by ALS conditions occurs as early as embryonic development.

**Figure 5.**
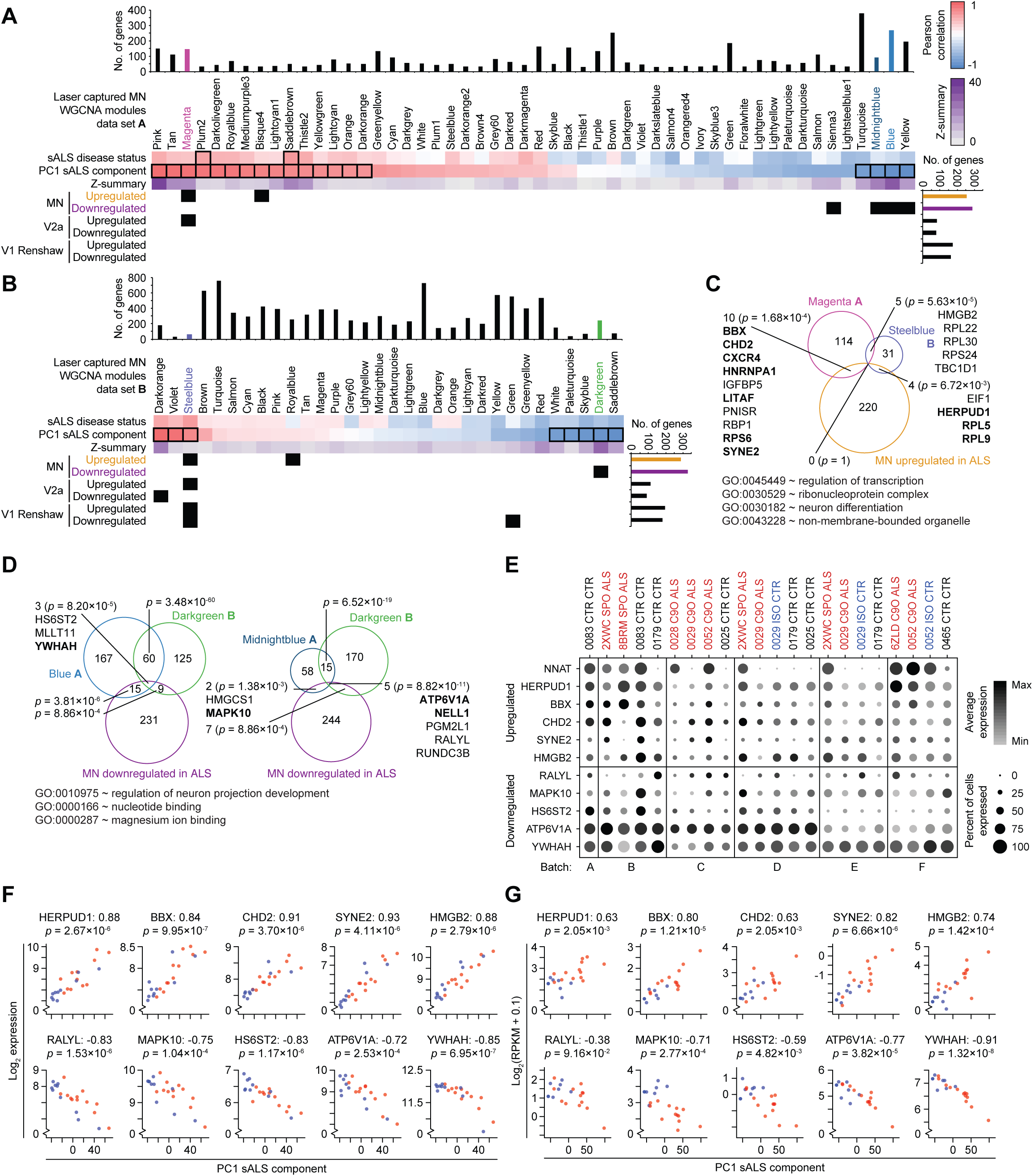
ALS iPSC-MN cultures exhibit transcriptional changes detectable in postmortem ALS spinal MNs. A. Hypergeometric test or each of 52 modules defined from WGCNA on LC MNs (data set A) from postmortem sporadic ALS (sALS) and non-ALS subjects (Ho et al., 2016) to detect enrichment for ALS dysregulated genes from each of the neuronal subtype categories identified in Figure 4C. Upper panel: WGCNA modules and the sample traits (sALS disease status and PC1 sALS component) with which they are significantly associated outlined in black rectangles (Pearson correlation between module eigengene and sample trait, Bonferroni-adjusted *P*-value < 0.01). Z-summary value for each module measures the extent of module preservation between data set A and B (Krach et al., 2018) (Figure 5B). Bar graphs above and to the right indicate the number of genes represented in data set A and our scRs data set, respectively. A matrix of *P*-values from hypergeometric tests performed for each module to neuronal subtype category comparison were adjusted by the Benjamini–Hochberg method, and subsequent *P*-values < 0.05 are marked as black squares in the matrix. B. As in A, except for 32 modules define from data set B (Krach et al., 2018). C. Euler diagram of intersecting gene sets among the Magenta module from data set A, the Steelblue module from data set B, and the MN upregulated in ALS genes. *P*-values indicate the likelihood of multi-set intersections using SuperExactTest (Wang et al., 2015). Genes are listed for overlapping sets, bolded genes are associated with the representative GO terms listed below. D. As in C, except among the Blue module from data set A (left), the Midnightblue module from data set A (right), the Darkgreen module from data set B, and the MN downregulated in ALS genes. E. Split dot plots indicate percent of MNs within each sample that express a non-zero value of each gene and also indicate the average gene expression among MNs within each sample that express non-zero values of that gene. F. Scatterplots depict gene expression in each LC MN sample from data set A (y-axis) against the coordinate for that sample along the first principal component (PC1 sALS component, x-axis). Prior to performing PCA on the expression data set, the ten genes shown in this panel were removed from the expression matrix to eliminate autocorrelation. sALS samples are colored red, and control samples are colored blue. The Spearman correlation between expression and PC1 coordinate is indicated next to the gene symbol, and the nominal *P*-value of the correlation is indicated below the gene symbol. G. As in F, except applied to data set B. See also Figure S6.

A closer examination of upregulated genes overlapping among the Magenta module in data set A, the Steelblue module in data set B, and MNs highlighted genes previously implicated in ALS and other motor neuropathies, and the overlapping genes and GO terms enriched among them are consistent with reports of disrupted mRNA and protein processing pathways (Deshaies et al., 2018; Kim et al., 2013, 2008; Montibeller and de Belleroche, 2018) (Figure 5C). Similarly, examination of downregulated genes overlapping among the Blue and Midnightblue modules in data set A, the Darkgreen module in data set B, and MNs highlighted genes previously implicated in ALS (Lederer et al., 2007; Saris et al., 2009; Umahara et al., 2016) (Figure 5D). The GO term regulation of neuronal projection development was significantly represented among the overlapping, downregulated genes (Figure 5D), consistent with recent models suggesting that deficiencies in maintaining axonal projections may underlie ALS (Klim et al., 2019; Melamed et al., 2019).

Auditing the expression of these overlapping genes in MNs demonstrated their dysregulation in ALS conditions as measured by average as well as percent expression (Figure 5E). Neuronatin (NNAT), which has been associated with neuronal development as well as degeneration (Joseph, 2014), was found to be upregulated in ALS MNs in the greatest number of ALS to control comparisons (Table S6A) while not observed as belonging to any modules significantly associated with sporadic ALS in postmortem data sets. Auditing the expression of ten overlapping genes in LC MNs from data sets A and B demonstrated high correlation between their expression and the first principal component that distinguishes sporadic ALS from control conditions (Figures 5F and 5G), further supporting the efficacy and fidelity of our discovery approach. A deeper investigation into the module genes disrupted in sporadic ALS conditions revealed a significant enrichment of module genes previously associated with spinal MN maturation and aging (Ho et al., 2016) (Figure 6).

**Figure 6.**
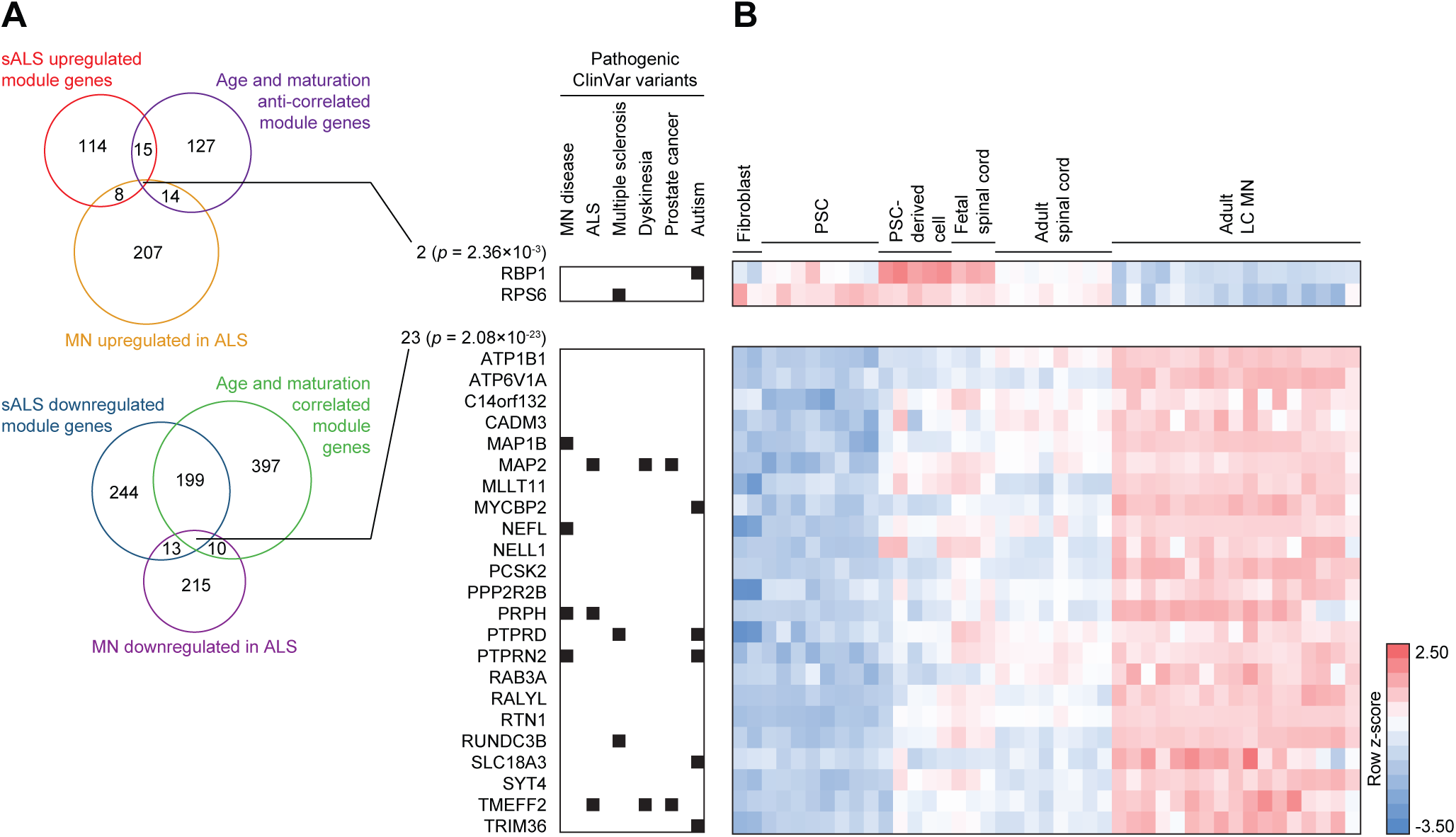
ALS iPSC-MN neuron cultures exhibit early transcriptional changes counteracting homeostatic maturation and aging. A. Euler diagram of intersecting gene sets among the Magenta, Blue, Midnightblue, or Yellow modules from data set A, the MN dysregulated genes in ALS identified in Figure 4C, and genes assigned to modules that significantly correlate with MN maturation and aging (Ho et al., 2016). The number of genes within each set is indicated. *P*-values indicate the likelihood of multi-set intersections using SuperExactTest (Wang et al., 2015). Genes with pathogenic variants in the ClinVar database are indicated on the right. B. Heatmap of genes listed in A demonstrating expression kinetics as tissues progress from embryonic, fetal, and adult spinal cord tissues (Ho et al., 2016). Pluripotent stem cells (PSCs) include embryonic stem cells and iPSCs.

### Predictive ALS markers are detectable in iPSC-MNs

While pooling of sparsely dysregulated genes in iPSC-MNs enriched for concordantly dysregulated genes in postmortem MNs, their average and percent expression varied considerably across subject lines (Figure 5E). This demonstrated a challenge in discovering consistently dysregulated genes by applying a significance threshold on a sample to sample basis across many scRs samples. We therefore took an alternative approach to discover genes that are consistently altered in iPSC-MNs from ALS subjects. We considered a combined expression score that reflected the average and percent expression for each gene in the MN populations at day 18 per subject (n = 22) in the scRs data set (see Methods). We then performed t-tests comparing the combined expression scores between all ALS and control and isogenic samples and ranked them by increasing nominal p-values. Among the top 20 ranking genes, we found six genes were concordantly downregulated in ALS conditions in data sets A and B, and they exhibited more uniform downregulation in ALS iPSC-MNs compared to controls (Figure 7A). We found no genes concordantly upregulated in all three data sets. Observing the expression kinetics of these genes over the course of embryonic, fetal, and adult spinal cord tissues (Ho et al., 2016) showed that some positively correlated with spinal MN maturation (ADCYAP1, ELAVL3, and NUAK1), DNMT3B anti-correlated with spinal MN maturation, and NDUFAF5 as well as NNAT were upregulated during fetal spinal cord stages (Figure 7B).

**Figure 7.**
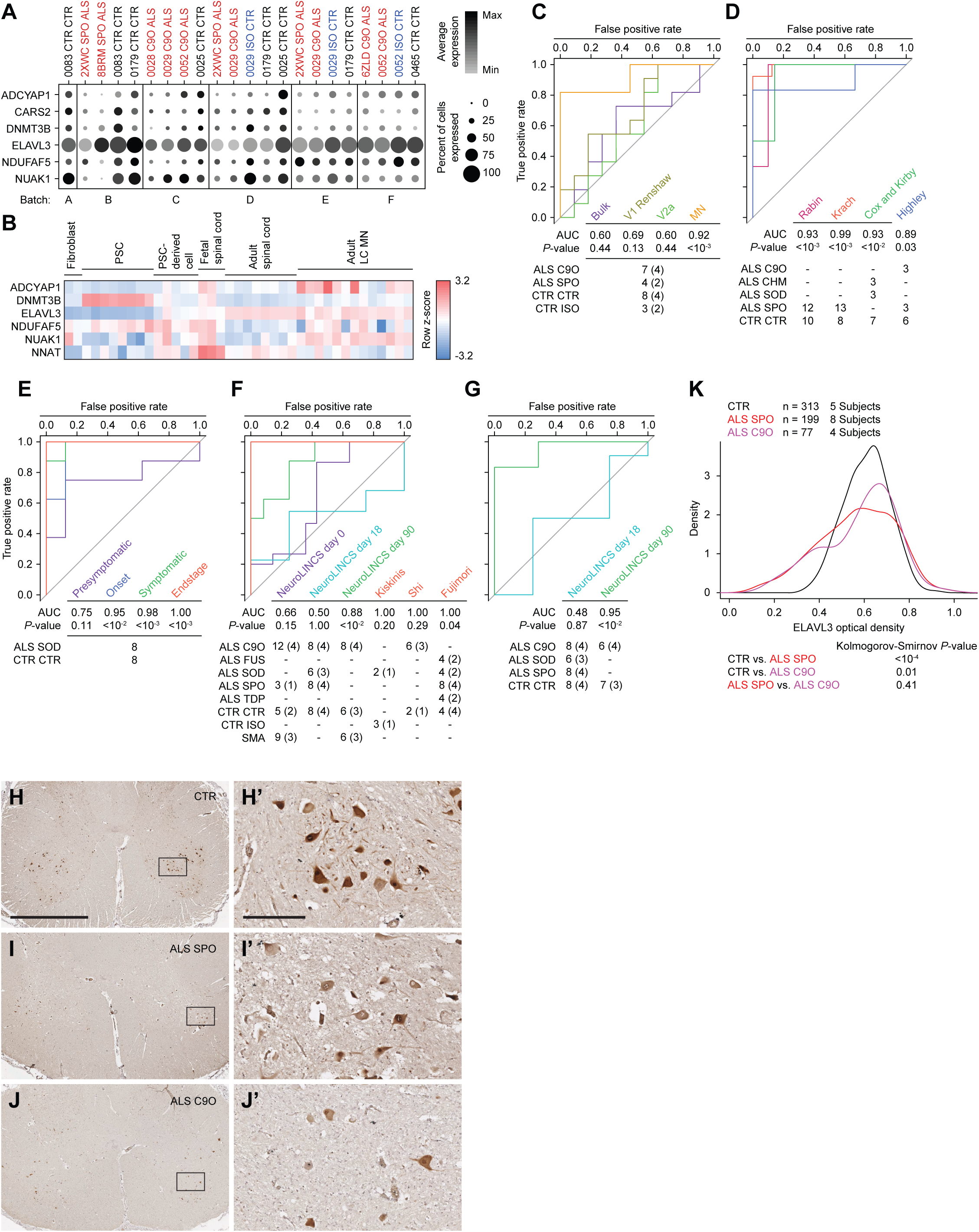
Single-cell analysis reveals predictive ALS marker genes. A. Split dot plots indicate percent of MNs within each sample that express a non-zero value of each of six predictive marker genes. Plots also indicate the average gene expression among MNs within each sample that express non-zero values of that gene. B. Heatmap of five genes listed in A, along with NNAT demonstrating expression kinetics as tissues progress from embryonic, fetal, and adult spinal cord tissues (Ho et al., 2016). CARS2 was not annotated in this data set. Pluripotent stem cells (PSCs) include embryonic stem cells and iPSCs. C - G. Receiver-operator characteristic (ROC) analysis performed on several data sets classifying samples in each data set as ALS or non-ALS, based on sample coordinates along the first, second, or both principal components using six predictive marker genes. See methods for calculations. P-values for each area under the curve (AUC) are calculated using the Mann-Whitney U test to determine whether the AUC differs significantly from 0.5 (diagonal grey line), which indicates an uninformative test. The number of samples for each ALS and non-ALS cases used in the analysis, along with their genotypes are indicated. The number of genetically distinct subjects included in the analysis are indicated in parentheses. Familial ALS subjects with pathogenic variants in C9orf72 (C9O), CHMP2B (CHM), FUS, SOD1 (SOD), TARDBP (TDP). Sporadic ALS subjects with no known pathogenic variants (SPO). Non-ALS control subjects (CTR). C9orf72 or SOD1 subject lines in which the pathogenic variant has been genome edited (ISO). Spinal muscular atrophy subjects classified as non-ALS (SMA). C. ROC analysis performed on subpopulation data generated from scRs data set based on a combined metric of average and percent expression for the six predictive marker genes. D. ROC analysis performed on previously published LC MN expression data sets (Cox et al., 2010; Highley et al., 2014; Kirby et al., 2011; Krach et al., 2018; Rabin et al., 2010). E. ROC analysis performed on mouse LCM MNs (Nardo et al., 2013). F. ROC analysis performed on bulk RNA-seq data sets generated by the NeuroLINCS Consortium for undifferentiated iPSCs (day 0), two MN differentiation protocols at days 18 and 90, and iPSC-MNs from Fujimori et al., 2018; Kiskinis et al., and 2014; Shi et al., 2018. G. ROC analysis performed on bulk proteomics data sets generated by the NeuroLINCS Consortium for two MN differentiation protocols at days 18 and 90. ELAVL3 protein expression is used as the prediction. H - J. Representative images of lumbar spinal cord sections from control, sporadic, and C9orf72 ALS subjects immunostained for ELAVL3 and counterstained with hematoxylin. Scale bar, 2.5 mm for H, I, and J; 250 um for inset images H’, I’, and J’. K. Comparative distributions of ELAVL3 optical densities. See also Figure S7.

The classification accuracy of ALS cases versus controls, as measured by the area under the curve (AUC), using PCA based on these six genes was significant in the MN population (Figures 7C and S7A). However, classification accuracy of ALS cases versus controls was not significant in V1 Renshaw, V2a, or by using bulk expression data (Figures 7C and S7A). The classification of sporadic ALS cases versus control postmortem adult spinal MNs in data set A (Rabin et al., 2010) and data set B (Krach et al., 2018) was also significant (Figure 7D and S7B). These results were expected, because the classifier genes were defined by these data sets. However, validating the accuracy of these six genes in classifying external test data sets would underscore their predictive power. In separate test data sets of postmortem adult spinal MNs from familial and sporadic ALS cases, which include variants in C9orf72, CHMP2B, and SOD1 (Cox et al., 2010; Highley et al., 2014; Kirby et al., 2011), classification using these genes significantly distinguished ALS from control subjects (Figures 7D and S7B). Additionally, in a disease progression study of SOD1G93A transgenic mouse spinal MNs (Nardo et al., 2013), classification of ALS versus control conditions based on these genes increased accuracy as mice progressed to the end stage of disease (Figures 7E and S7C). Interestingly, focusing analysis on these genes from the NeuroLINCS Consortium (Keenan et al., 2018) bulk RNA-seq data set analyzing undifferentiated human iPSCs and iPSCs differentiated into MN cultures over 18 days demonstrated that ALS could not accurately be distinguished from control conditions (Figure 7F and S7D). However, using a different and longer MN differentiation protocol (Sareen et al., 2013), where MN cultures were extended up to 90 days and again profiled by bulk RNA-seq, analysis of these six genes demonstrated a significant accuracy in classifying ALS cases from control as well as from spinal muscular atrophy cases (Figures 7F and S7D). Additionally, these signature genes could distinguish SOD1A4V ALS patient samples from zinc-finger nuclease corrected isogenic samples in iPSC-derived, HB9-RFP positive MNs at 39 days of differentiation (Kiskinis et al., 2014) (Figures 7F and S7D). Similarly, these genes also distinguished C9orf72 ALS patient derived, HB9-RFP positive MNs from control samples, and further distinguished isogenic control samples in which one or two copies of the C9orf72 HRE were targeted into the genome with CRISPR-Cas9 (Shi et al., 2018) (Figure 7F). Finally, this panel of genes distinguished control subject iPSC-MN cultures from sporadic and familial ALS subjects, including those with variants in FUS, SOD1, and TARDBP (Fujimori et al., 2018) (Figure 7F). Among the six genes quantifiable by RNA in all expression data sets tested, ELAVL3 was the only gene quantifiable as a protein when analyzing the NeuroLINCS proteomics data sets at 18 and 90 days of differentiation. Classification of iPSC-MN cultures based solely on ELAVL3 protein expression demonstrated that it was an accurate classifier for ALS versus control MN cultures only in extended cultures at 90 days, where it was also decreased in ALS conditions (Figure 7G). However, analysis of ELAVL3 RNA alone demonstrated less overall accuracy when compared to joint RNA analysis of all six genes (Figures S7E-H). Lastly, the decline in ELAVL3 protein per MN was also detectable in postmortem spinal cords in both sporadic and C9orf72 ALS cases versus control (Figures 7H-7K). Altogether these data reveal that despite globally resembling in vivo fetal tissue, single-cell analysis of iPSC-MNs can model early, common signatures of familial and sporadic ALS that persist into the end stages of disease.

## Discussion

Recent scRs studies have characterized diverse neuronal populations in vivo mouse spinal cords (Delile et al., 2019; Sathyamurthy et al., 2018). However, this technology has not been used to rebuild the spinal cord from complex mixtures of cells in cultures differentiated from iPSCs. Our approach described here is ideally suited to achieve this goal and demonstrates initial steps toward building a iPSC-based cellular atlas of the developing human spinal cord to provide an anatomical context for human embryonic development as well as disease modeling.

### Overcoming experimental challenges to single-cell profiling iPSC models to cope with technical and biological variability

As variable molecular readouts caused by genetic background is becoming increasingly acknowledged by experimentalist in human iPSC disease modeling, experimental design must account for the genetic backgrounds of several individual subjects as well as isogenic controls in order to isolate reproducible disease-related effects (Fujimori et al., 2018; Kiskinis et al., 2014; Shi et al., 2018). In line with this outlook, we incorporated iPSC lines from several ALS and control subjects into this study, and repeatedly assayed MN differentiations with the goal of detecting reproducible transcriptional signatures in distinct cellular subpopulations. However, repeated experimental sampling presented the challenge of coping with batch effects, which in the process of scRs analysis, severely affects global clustering approaches towards cell type annotation such as Louvain community detection and tSNE dimensionality reduction (Hicks et al., 2018; Luecken and Theis, 2019). We alleviated these effects on these unsupervised learning methods through MultiCCA (Butler et al., 2018). In addition, we referenced decades’ worth of developmental genetic literature to supervise cell type annotation using a relatively small number of key developmental genes (Alaynick et al., 2011; Lu et al., 2015). This complementary approach was more refractory to batch effects, and when combined with batch-corrected global clustering, enabled us to confidently annotate cell identities.

Having reliably annotated cells uniformly across experimental batches, the next challenge was to find reproducibly dysregulated genes between ALS and control conditions within each subpopulation. Despite increasing sample size and replicates, the sparse nature of scRs yielded low reproducibility of dysregulated genes. This was especially evident in CRISPR-Cas9-corrected isogenic comparisions, despite the observations that genetic removal of the C9orf72 HRE universally corrected RNA and dipeptide phenotypes. This suggests that even in conditions that rule out genetic variability, sensitivities to technical noise arising from experimental platform or batch variation remain. To overcome the sparsity of reproducibly dysregulated genes, we relaxed the threshold in cataloguing upregulated and downregulated genes by counting instances where genes are detected in any comparison more than once and then subsequently pooling them together. Interestingly, genes defined as dysregulated by this criteria in C9orf72 HRE ALS conditions significantly overlapped with those determined in sporadic ALS conditions. This motivated us to analyze C9orf72 HRE and sporadic conditions together to jointly discover genes, co-expression networks, and pathways that are convergently disrupted in both conditions. As an alternative to selecting genes based on significance thresholds, we also applied a ranked metric to discover consistently dysregulated genes across ALS and control conditions that did not meet significance. This approach revealed a small number of genes that were consistently downregulated in ALS in the scRs data and additionally served as effective predictors of ALS cases. Importantly, we validated this methodology by testing our discoveries against external data sets such as RNA-seq data from postmortem LC MNs or other iPSC-based or animal models of familial and sporadic ALS. We validated both cell annotation as well as ALS signatures, as further described below.

### Single-cell resolution of rostral, caudal, hindbrain, and spinal cord identity

The colinear expression of HOX genes during embryonic development spatially defines rostral to caudal body segments and downstream neural fate (Di Bonito et al., 2013; Lippmann et al., 2015; Philippidou and Dasen, 2013). HOX gene expression from bulk sample preparations of iPSC-differentiated neurectoderm cultures suggested that more caudal identities are present (Lippmann et al., 2015). Our scRs analysis of day 18 cultures, which we demonstrated to be globally similar to approximately 42 days of in vivo embryonic development, enabled the resolution of rare cells that have activated expression of caudal identity-encoding HOX genes. This revealed the majority of cells were similar to hindbrain and cervical spinal cord regions. This anatomical classification lays the foundation for future work with human iPSC models to investigate instrinsically different physiologies between rostrocaudal regions that contain cranial and phrenic MNs, respectively.

iPSC-based models are further challenged in that the cells differentiated in vitro do not fully recapitulate the distinctive identities of cells formed in vivo. Therefore, cellular identities are less resolved by conventional dimensional reduction and classification based on a small set of markers, even after batch correction. By using a unit expression matrix of genes assiduously delineated by Thomas Jessell and others to define cell types in the developing chick and rodent spinal cords (Alaynick et al., 2011; Delile et al., 2019; Lu et al., 2015), we assigned identities based on significant Pearson correlation to the profiles in this expression matrix, which accounts for both the presence and absence of key marker gene expression in specific cell types. Together this combined classification method dissected out specific neuronal and progenitor populations from complex, in vitro tissue cultures to rebuild sections of in vivo hindbrain and spinal cord.

### Single-cell analysis enables attribution of early and convergent ALS signatures to MNs

The most recent iPSC-based transcriptomic reports performed RNA profiling at time points during differentiation concomitant with various observed ALS phenotypes which include nuclear RNA foci (Sareen et al., 2013), decreased neurite length (Fujimori et al., 2018), reduced neurite repair after injury (Klim et al., 2019; Melamed et al., 2019), and MN death (Kiskinis et al., 2014; Shi et al., 2018). Several of these protocols differentiated iPSCs for over 30 days, and many required a relatively prolonged maturation phase, the presence of glia, and additional stressor conditions in order to yield a disease phenotype. Thus, it is unclear whether the transcriptional events observed precede the disease phenotypes, are concomitant, or are immediate consequences of other prior events. We elected to profile transcription in postmitotic MNs at an earlier point, day 18 at which their identity was established and in the absence of glial cells. This event was demonstrated as early as day 14 of differentiation (Maury et al., 2015). Our approach satisfied two objectives. One was to capture a transcriptional signature as early as possible, prior to the manifestation of disease phenotypes ranging from neurite repair to overt cell death. The other was to reduce heterogeneity across subject lines and experimental batches that could be augmented by a longer time of differentiation in culture. Within this early developmental time point, we detected signatures in ALS conditions prior to disease phenotypes, suggesting that these transcriptomic events precede and are potentially causative of later phenotypes.

The ability to leverage expression information on a panel of MN marker genes bolstered our validation that iPSC-derived MNs globally resemble adult in vivo MNs more than any other cell types concommittantly arising in culture. Additionally, single-cell profiling of all cells present in the differentiation cultures enabled us to account for the direct contribution of each cell type to the signals present in bulk RNA-seq. The dysregulation of gene expression we discovered in V1 Renshaw interneurons supports previous work that have proposed interneuron dysfunction and degeneration as a cause of MN changes in ALS (Chang and Martin, 2009; Maekawa et al., 2004; Martin and Chang, 2012; Morrison et al., 1998; Stephens et al., 2006; van Zundert et al., 2008). Future work exploring the contributions of interneurons to ALS in iPSC-MN cultures may provide more mechanistic insights into this model of disease.

Of note, recent literature has characterized convergent and divergent aspects of ALS pathways and mechanisms across subject lines and subsets of ALS conditions, often delineated by genotype (including familial ALS mutations in C9orf72, SOD1, hnRNPA1, TARDBP, FUS, and CHMP2B as well as sporadic ALS), site of symptom onset, disease progression, or treatment response (Chen et al., 2014; Fujimori et al., 2018; Kiskinis et al., 2014; Klim et al., 2019; Melamed et al., 2019; Sareen et al., 2013; Shi et al., 2018). We noted that bulk transcriptomic analysis may contribute to discrepancies in pathway discoveries, and thus sought to characterize these commonalities and distinctions using higher resolution, single-cell analysis. These changes were observed specifically in MNs and can be used to establish whether the MNs were from a control or ALS subject. This ability to determine disease phenotype with greater accuracy means that an at-risk ALS subject could have iPSCs generated and differentiated into MNs. Following scRs, it would then be possible to predict whether the subject would go on to get ALS with some fidelity. As more samples are run through this pipeline it will be possible to further refine the the subtype of ALS through correlative studies. Thus, the six genes we have defined based on our iPSC-MNs and postmortem laser-captured MNs can be used as early biomarkers across a broad range of familial and sporadic ALS cases, and they can be further expanded upon through the discovery approach we have described here to improve accuracy.

### Single-cell analysis of ALS iPSC-MNs highlights putatively causal genes and pathways

Despite their global resemblance to fetal tissue, iPSC-MNs from ALS subjects exhibited concordant signatures detectable in MNs from postmortem ALS subjects. While these observations highlight the utility in discovering effective disease biomarkers, a more potent utility would be to discover disease drivers. In order to determine if these early changes in gene expression are causative of ALS, the complete course of disease should be recapitulated in iPSC-based models to ascertain this causality. We have previously demonstrated that ALS preferentially disrupts homeostatic pathways enacted during neuronal maturation and aging contexts (Ho et al., 2016). By intersecting the set of genes disrupted in MNs with our previously characterized gene co-expression modules involved with MN maturation, aging, and ALS, we revealed that pathways associated with neurite structure and synaptic vesicle trafficking, which increase with maturation and aging, are downregulated in ALS subject cells, and these changes are already detectable in iPSC-MNs. This suggests that early deficiencies in activating these homeostatic pathways may prime the MNs to a disease state upon maturation and aging. We posit that strategies that accelerate maturation and aging gene expression programs in patient iPSC models can enact a state where the ALS-driven deficiencies, which possibly stem from these early gene expression changes, can be observed and therapeutically treated (Studer et al., 2015).

The most notable among the marker genes we have highlighted is ELAVL3, whose dysregulation was detectable in various ALS conditions during early, late, and end stages of diseased MNs. It is a neuron-specific RNA-binding protein whose expression starts embryonically and persists into adulthood (Okano and Darnell, 1997). Previous studies have noted disruption of ELAVL3 in ALS conditions (Colombrita et al., 2015; Klim et al., 2019), and because it regulates RNA processing, splicing, alternative poly-adenylation, and stabilizes translational control (Grassi et al., 2018), future studies investigating functional disruption of ELAVL3 may address whether it is an upstream pathway to MN degeneration shared among familial and sporadic ALS cases.

Finally, future iPSC-based studies that distinguish bulbar from spinal onset ALS patients can build upon the data reported here to help correlate region of onset in the patient with the pathology in specific MNs associated with those regions. Equally important, our anatomical assessment of iPSC-MN models establishes a cellular and molecular framework to address how MN degeneration and paralysis spreads throughout the body of ALS patients, mechanisms which are of great interest to develop accurate prognostic assessments or interventional therapies (Turner et al., 2010). While the fidelity of our iPSC-MNs to in vivo MNs was based on pooled LC MNs, recent advances in single nuclei RNA-seq of human postmortem tissues of the central nervous system (Gaublomme et al., 2019; Mathys et al., 2019) will provide an expanded resolution of cellular and disease signatures with which our data can be reconciled. This comparison will enable better interpretation of molecular signatures and cellular compositions as they arise in early stages of ALS and progress into the end stages of ALS, thus enabling a better understanding of disease etiology. Finally, the analysis reported here provides a methodological resource for iPSC-based disease models of not only ALS, but also for several other late onset diseases standing to benefit from single-cell resolved investigations.

## Methods

### EXPERIMENTAL MODEL AND SUBJECT DETAILS

All human iPSC lines are banked and available through the Cedars-Sinai Biomanufacturing Center. Cell lines were routinely characterized for mycoplasma, Alkaline Phosphatase staining, immunostaining for pluripotency markers, karyotypes by G-banding, PluriTest, Trilineage Differentiation Potential (assessed via TaqMan hPSC Scorecard Assay), and Cell Line Authentication (assessed via STR Analysis) to match primary donor tissue. Relevant clinical and experimental data about the iPSC donor subjects (e.g. age, sex, tissue source) are presented in Table S1 and in the Key Resources Table. All protocols were performed in accordance with the Institutional Review Board guidelines at Cedars-Sinai Medical Center under the auspices of IRB-SCRO Protocol 21505.

## METHODS DETAILS

### Culture of human iPSCs

All iPSC lines were maintained in complete mTeSR1 growth medium on Growth Factor Reduced Matrigel and passaged every seven days using the StemPro EZ Passaging Tool or Versene and typically split between 1:4 and 1:9 ratios.

### Genome editing of C9orf72 repeats in iPSCs

CRISPR guides were designed to target regions immediately 5’ and 3’ of the C9orf72 hexanucleotide repeat expansion using the Zhang lab CRISPR design tool (Shi et al., 2018). Guides were cloned into pSpCas9(BB)-2A-GFP (PX458) plasmids, (gift from Feng Zhang, Addgene plasmid #48138. Each iPSC line was transfected with both 5’ and 3’ targeting plasmids using the Neon Electroporation System (Thermo Fisher). After 48 hours, iPSCs were dissociated and flow sorted by GFP fluorescence to isolate successfully transfected cells. These cells were plated, cultured for 1 week, passaged, and allowed to grow to confluency. Cells were then subcloned as following: iPSCs were Accutase-dissociated into single cells and replated sparsely at 30,000 cells/10 cm dish. Rock inhibitor (Y-27632) was included for 24 hours after plating to promote iPSC survival. Once individual cells formed small colonies, pipette tips were used to manually transfer subclones from the 10 cm dish into individual wells of a 96 well plate. These subclones were passaged with Versene into two 96 well plates, one for further propagation and one for gDNA extraction and sequencing. Sanger sequencing was used to identify subclones that had modified genetic sequences around the C9orf72 locus. PCR products from these subclones were then TOPO-cloned (Invitrogen) and Sanger sequenced to determine the sequences of each allele. Subclones lacking the C9orf72 HRE were expanded and characterized.

### Repeat primed PCR assay for HRE

PCR products were amplified using FastStart Master Mix (Roche) and 1X betaine using the following cycling conditions: 1x 95°C for 15 min, 2x 94°C for 1 min -> 70°C for 1 min -> 72°C for 3 min, 3x 94°C for 1 min -> 68°C for 1 min -> 72°C for 3 min, 4x 94°C for 1 min -> 66°C for 1 min -> 72°C for 3 min, 5x 94°C for 1 min -> 64°C for 1 min -> 72°C for 3 min, 6x 94°C for 1 min -> 62°C for 1 min -> 72°C for 3 min, 7x 94°C for 1 min -> 60°C for 1 min -> 72°C for 3 min, 8x 94°C for 1 min -> 58°C for 1 min -> 72°C for 3 min, 5x 94°C for 1 min -> 56°C for 1 min -> 72°C for 3 min, and 1x 72°C for 10 min. Samples were then sent to Genewiz for sequencing.

### Sanger sequencing of C9orf72 locus

The C9orf72 locus was PCR-amplified using PrimeStar Polymerase with 1X betaine. To determine the sequence of each allele, PCR products were cloned using the TOPO Cloning Kit for Sequencing (Invitrogen). Plasmids were used to transform TOP10 competent bacteria, which were plated on agar dishes containing ampicillin and incubated at 37°C overnight. Bacterial plates were sent for direct colony sequencing at Genewiz.

### Karyotype

G-Band Karyotyping was performed by the Cedars-Sinai Biomanufacturing Center.

### Differentiating iPSC-MN cultures

For Figures S2A-E, iPSC-MNs as previously described (Yang et al., 2013). In brief, iPSCs were dissociated into single cells, cultured in Neural Induction Media (NIM) consisting of Neurobasal (Gibco), 1.1 µM ascorbic acid (Sigma), 1% non-essential amino acids (Gibco), 1% GlutaMax (Gibco), 2% B27 (Gibco), 0.16% D-glucose solution, and 1% penicillin-streptomyosin-amphotericin solution. 10 µM Y-27632 ROCK inhibitor (Sigma) was included in the media or the first 48 hours to improve survival of iPSCs following dissociation. On days 1-4, NIM was supplemented with 10 µM SB431542 (StemGent), 1µM Dorsomorphin (Sigma), and 10 ng/mL bFGF (PeproTech). This media was changed every other day, and bFGF was replenished daily. On day 5, cells were cultured in NIM supplemented with SB431542, Dorsomorphin, 10 ng/mL BDNF (R&D), and 1 µM retinoic acid (Sigma). On days 7 and 9, the media was changed to NIM with BDNF, retinoic acid, and 1 µM smoothen agonist (Sigma). The cells were densely plated onto poly-ornithine/laminin coated dishes and cultured in the same media on day 11. This media was further supplemented with 2 µM DAPT (Cayman Chemicals) on days 13-20, with media changes every 2-3 days. On days 20-30, cells were fed every 2-3 days with NIM containing BDNF, smoothen agonist, 1 µM retinoic acid, and 2 µM Ara-C (Sigma). Cells were then gently dissociated using papain (Worthington), plated on poly-ornithine/laminin coated dishes, and cultured in NIM with the addition of 1% N2 supplement (Gibco), 4 µM Ara-C, and 40 ng/mL each of growth factors BDNF and GDNF (PeproTech). For Figures S2F and S2G, iPSC-MN cultures were differentiated using the 18 day protocol as described below for polyGP ELISA.

For the 18 day iPSC-MN differentiation, mTeSR1 was removed from iPSCs at 30-40% confluency and replaced with Stage 1 media (1:1 mixture of Iscove’s Modified Dulbecco’s Medium (IMDM):F12 basal media supplemented with 1% non-essential amino acids (NEAA), 2% B27, 1% N2, 1% Penicillin-Streptomycin-Amphotericin B solution (PSA), 0.2 μM LDN193189 (Selleck), 10 μM SB431542, and 3 μM CHIR99021(Xcess Biosciences)) for six days with daily media changes. The cells were then Accutase-treated to single-cell suspension and centrifuged in 50 ml conical tubes, resuspended in Stage 2 media (Stage 1 media further supplemented with 0.1 μM all-trans retinoic acid (Stemgent) and 1 μM Sonic hedgehog agonist (SAG) (Cayman Chemicals)), and plated onto Matrigel-coated plates or coverslips. Stage 2 media was changed every two days until day 12, when Stage 3 media (1:1 mixture of IMDM:F12 basal media supplemented with 1% non-essential amino acids (NEAA), 2% B27, 1% N2, 1% PSA, 0.1 μM Compound E (Calbiochem), 2.5 μM DAPT, 0.1 μM dibutyryl cyclic adenosine monophosphate (db-cAMP), 0.5 μM all-trans retinoic acid, 0.1 μM SAG, 200 ng/ml ascorbic acid, 10 ng/ml BDNF, and 10 ng/ml GDNF) was then used to feed cells every two days until day 18, when cultures were analyzed.

The 90 day iPSC-MN differentiation was performed as previously described (Ho et al., 2016). In brief, 80% confluent iPSC cultures were Accutase-treated into single cells suspension and centrifuged in 384-well Matrigel coated PCR plates. The cells were maintained in Neural Differentiation Media (NDM): IMDM/F12 supplemented with 2% B27-vitamin A, 1% N2, 1% NEAA, 0.2 µM LDN193189, and 10 µM SB431542. On day two, neural aggregates were collected and transferred into Poly-Hema coated T75 flasks and the aggregates were cultured for three more days in NDM. On day seven, aggregates were collected and transferred onto poly-ornithine/laminin coated wells with fresh NDM. After five days, cells were cultured in MN Specification Media: NDM supplemented with 0.25 µM all-trans retinoic acid, 1 µM purmorphamine, 1 µM db-cAMP, 200 ng/mL ascorbic acid, 20 ng/mL BDNF, and 20 ng/mL GDNF. Once rosettes were observed, they were collected with STEMdiff Neural Rosette Selection Reagent and cultured in MN Precursor Expansion Media: NDM supplemented with 0.1 µM all-trans retinoic acid, 1 µM purmorphamine (Millipore), 100 ng/mL EGF, and 100 ng/mL FGF2. After day 26, the iPSC-MN precursor spheres (iMPS) are expanded by using a chopping method every seven to ten days. The iMPS are matured into MNs for 21 days in MN Maturation Media: Neurobasal supplemented with 1% NEAA, 0.5% Glutamax, 1% N2, 10 ng/ml BDNF, 10 ng/ml GDNF, 200 ng/ml ascorbic acid, 1 µM db-cAMP and 0.1 µM all-trans retinoic acid.

### Immunofluorescent staining, imaging, and quantification of iPSC-MN cultures

iPSC-MNs were fixed in 4% paraformaldehyde, rinsed with PBS, incubated in 0.5% Triton-X in PBS, rinsed with 0.2% Tween-20 in PBS, incubated in blocking solution (5% normal donkey serum and 0.2% Tween-20 in PBS). Primary antibody solution in blocking solution containing various combinations of goat polyclonal IgG anti-Human ISL1 (1:200) (R&D Systems AF1837, RRID: AB_2126324), mouse monoclonal IgG1 anti-NF-H (SMI-32) (1:200) (BioLegend 801701, RRID: AB_2564642), goat polyclonal anti-ChAT (1:200) (Millipore AB144P, RRID: AB_2079751), rabbit polyclonal IgG anti-PHOX2B (1:200) (GeneTex GTX109677, RRID: AB_1951223), mouse monoclonal IgG2b, rabbit polyclonal IgG anti-CHX10 (VSX2) (1:200) (Novus NBP1-84476, RRID: AB_11022841), and rabbit polyclonal IgG anti-SOX1 [EPR4766] 1:200) (GeneTex GTX62974) were incubated, rinsed with 0.2% Tween-20 in PBS, and incubated in species-specific Alexa-fluor secondary antibodies (1:2,000), and rinsed with 0.2% Tween-20 in PBS with DAPI staining. Fluorescent images were acquired using ImageXpress Micro XLS system (Molecular Devices) at 10X magnification. For a complete analysis, total 9 sites per well were captured. The captured images were quantified for the cellular population using MetaXpress software (Molecular Devices).

### Quantification of C9orf72 transcript variants

RNA was extracted from iPSCs using the PureLink RNA mini Kit (Invitrogen) and reverse-transcribed into cDNA using the Promega Reverse Transcription System. Quantitative PCRs were conducted in triplicate using SYBR Green and primers amplifying all C9orf72 transcripts as well as specific transcript variants. PCR cycles consisted of the following steps: [1x 95°C for 10 min, 40x 95°C for 30 seconds -> 58°C for 60s, and 1x 72°C for 5 min].

### FISH of C9orf72 sense and antisense RNA foci and imaging

RNA FISH was performed as previously described in (Sareen et al., 2013). Briefly, cells were cultured on chamber slides (Lab-Tek II chamber slide system, Thermo Fischer Scientific, Cat #154917). Cells were then fixed in 4% paraformaldehyde, permeabilized with diethylpyrocarbonate (DEPC)-PBS/0.2% Triton X-100, and washed with (DEPC)-PBS. Cells were incubated with hybridization buffer containing 50% formamide, DEPC-2xSSC (300 mM sodium chloride, 30 mM sodium citrate, pH 7.0), 10% w/v dextran sulfate, and DEPC-50 mM sodium phosphate, pH 7.0 for 30 min at 66°C. This was followed by hybridization with 40 nM of a Locked Nucleic Acid probe for C9orf72 HREs in hybridization buffer for 3 hours at 66°C. Afterwards, the cells were rinsed once in DEPC-2xSSC/0.1% Tween-20 at room temperature and three times in DEPC-0.1xSSC at 65°C. The cells were then stained with DAPI, mounted using ProLong Gold antifade reagent, and analyzed with fluorescence microscopy.

### PolyGP Response

PolyGP in iPSC-MNs were measured blinded to C9orf72 HRE and disease status using a previously described sandwich immunoassay that utilizes Meso Scale Discovery electrochemiluminescence detection technology, and an affinity purified rabbit polyclonal polyGP antibody (Rb9259) as both capture and detection antibody (Gendron et al., 2015; Su et al., 2014).

### Single-cell RNA-seq of MN cultures

iPSC and iPSC-MN differentiation cultures were washed with PBS, incubated at 37°C with 0.25% Trypsin-EDTA between 5 and 20 minutes, and diluted with an equal volume of the complete culture media in which they were grown. After pelleting cells at 200 x g for five minutes at 4°C, cells were resuspended in PBS, observed for clumps, and further triturated with a fire polished glass pipet. The cell suspension was filtered through a Miltenyi 30 µm filter, counted on a hemocytometer, and the concentration was adjusted prior to loading onto the Illumina Bio-Rad ddSEQ System or 10X Genomics Chromium scRs platforms in accordance with the respective instructions for each kit for targeting approximately 1,000 cells per sample. Library preparation kits used were Illumina® Bio-Rad® SureCell™ WTA 3’ Library Prep Kit for the ddSEQ™ System and 10X Chromium Single-cell 3’ Library & Gel Bead Kit v2. Libraries were sequenced on Illumina NextSeq500 targeting 100,000 reads per cell. Raw sequencing reads were demultiplexed and processed to FASTQ using Illumina bcl2fastq. Sample reads were aligned to the transcriptome and uniquely mapped reads were counted and assigned to cell specific barcodes. For ddSEQ libraries, reads were aligned and demultiplexed to cell barcodes using Illumina Single-Cell RNA Seq BaseSpace Workflow (v1.0.0) with STAR Aligner (v2.5.2b) (Dobin et al., 2013) and hg19 reference genome. For 10X libraries, reads were aligned and demultiplexed using 10X Genomics Cell Ranger (v2.1.0) with STAR Aligner (v2.5.1) and GRCh38 reference genome. Ensembl gene IDs were annotated to HGNC symbols. In instances of multiple ENSG IDs mapping to unique HGNC symbols, the sum of unique molecular identifiers (UMIs) across ENSG IDs was calculated and used as the UMI for the unique HGNC symbol. The summarized UMI count tables for each experimental batch are deposited in GEO under accession number GSE138121.

### Immunohistochemistry and quantification of ELAVL3 in spinal cords

Human tissues were obtained using a short-postmortem interval acquisition protocol that followed HIPAA-compliant informed consent procedures and were approved by Institutional Review Board (Benaroya Research Institute, Seattle, WA IRB# 10058 and University of California San Diego, San Diego, CA IRB# 120056). For IHC, 8 sporadic ALS, 4 C9 ALS, and 5 control lumbar spinal cord sections were studied. Sections with 6 µm thickness were formalin-fixed and paraffin-embedded. On day one, sections were deparaffinized with Citrisolv (Fisher Scientific #04-355-121) and hydrated with different dilutions of alcohol. Endogenous peroxidase activity was quenched with 0.06% H2O2 for 15 min. Antigen retrieval was performed in a Antigen Unmasking Solution (Vector Laboratories #H-3301) in a pressure cooker for 20 min at a temperature of 120 °C. Following antigen retrieval, sections were permeabilized with 1% FBS (Atlanta Biologicals #511150) and 0.2% Triton X-100 in PBS for 15 min and then blocked with 2% FBS in PBS for 25 min. The sections were incubated overnight with the primary antibody, rabbit polyclonal ELAVL3, 1:1000, LSBio, Cat# LS-C408905. On the second day, after 60-min incubation with the secondary antibody (Immpress reagent kit, anti-Rabbit, Vector Laboratories #MP-7401) in room temperature, signals were detected using Immpact DAB (Vector Laboratories #SK-4105) for 5–10 min. Counterstaining was performed with hematoxylin (Fisher #HHS128). For IHC visualization, slides were scanned with Hamamatsu Nanozoomer 2.0HT Slide Scanner at 40X magnification. At least 6 motor neurons per spinal cord were evaluated, and across all samples totaled a combined number of 199 neurons from sporadic ALS subjects, 77 neurons from C9 ALS subjects, and 313 neurons from control subjects. Images were deconvoluted using Fiji ImageJ (Schindelin et al., 2012) and the optical density (OD) was measured for each neuron, where OD = log (max intensity/1/Mean intensity), where max intensity = 255 for 8-bit images.

### QUANTIFICATION AND STATISTICAL ANALYSIS

#### Pseudotime analysis

Monocle version 2.12.0 (Qiu et al., 2017) was used to perform pseudotime analysis of the 18 day differentiation time course. Genes with minimum average expression of 0.1 and detectable in at least 10 cells were filtered. Cells were further filtered for those whose total UMI count was within three standard deviations of the average log10 UMI across all time points. Tests each gene for differential expression as a function of the time course was calculated using the full generalized linear model, and genes with a q-value less than 0.1 from this test was filtered. These genes were used in dimensional reduction of the time course samples onto two components through Discriminative Dimensionality Reduction with Trees. All cells were ordered along this pseudotime trajectory, and expression of select genes were plotted against the cells ordered along this pseudotime.

Seurat Version 2.3.0 was used to process, normalize, cluster, and analyze scRs data for day 18 MN cultures. UMI count tables for each of the six experimental batches were each loaded as Seurat objects as well as cell barcodes and sample covariates for meta data. Genes with at least one UMI in at least one cell were kept. The percent of mitochondrial genes was calculated for each cell and stored as meta data. Z-scores were calculated for three columns in the meta data for each cell: nGene, nUMI, and percent mitochondrial genes. Cells were then filtered based on these z-scores; any cell that had a z-score greater than 3 or less than −3 (greater than 3 standard deviations away from the mean of that meta data) in any of the three columns were excluded from further analysis. Next, the global scaling normalization method normalizes the gene expression measurement for each cell by the total expression, multiplies this by a scale factor, and log transforms the result. The maximum UMI detected in the experimental batch was used as the scaling factor. Next, highly variable genes (HVGs) in the experimental batch were calculated. The mean expression for all detected (i.e. non-zero value) genes was calculated as well as the log transformed ratio of variance to mean expression (regarded as the dispersion). Genes were then binned into 20 intervals, and within each interval, the z-score for dispersion was calculated for each gene. This helps control for the relationship between variability and average expression. Genes with z-score for dispersion values greater than 2 were deemed to be HVGs. After all six experimental batches were processed as Seurat objects, samples were subsetted out of each Seurat object, totaling 22 samples. 279 HVGs were calculated in at least 11 of the 22 samples, and these were kept for subsequent dimensional reduction.

### Data set normalization, identity assignment, and clustering

Multiple Canonical Correlation Analysis (MultiCCA) was performed on the 22 samples to correct for experimental batch and platform effects. Up to 20 dimensions were evaluated, and the first 18 dimensions were determined to be used for subspace alignment. Prior to subspace alignment, cells whose expression profiles cannot be well-explained by low-dimensional CCA compared to low-dimensional PCA (less than a two-fold ratio) were removed. 17,531 cells remained. Subsequently, samples were aligned using dynamic time warping along the first 18 dimensions, and the resulting batch integrated Seurat object holding all 22 samples was used for downstream analysis. To determine the optimal number of communities to cluster, several resolution settings were tested using the FindClusters command in Seurat. The first 18 dimensions from the reduction through CCA were used, and 30 nearest neighbors were considered for each resolution setting. All other parameters were kept at default values. The original Louvain algorithm determined the modularity for each setting, and the maximum modularity observed after 100 iterations was recorded for each number of communities. A polynomial trendline was calculated, and the residuals for each setting greater than zero was considered to determine the optimal number of communities. Based on the independently optimized tSNE calculations and visualizations for 17,531 cells, a resolution setting of 0.125 yielding 4 communities was selected to proceed with downstream analysis. When projecting all cells on two two dimensional tSNE plots using the RunTSNE command, the same 18 dimensions were used as for the FindClusters command, and all other parameters were kept at default values. A perplexity setting of 100 was selected based on the visual concordance with the 4 communities determined.

To analyze only the postmitotic, neuronal subtypes from these 17,531 cells, we repeated the FindClusters command using a resolution parameter of 0.04, which detected 2 communities, and the postmitotic community containing 11,120 cells was subsetted into a new Seurat object. Once again, 22 samples were subsetted out of this Seurat object. HVGs were again calculated in each of the 22 samples using the same parameters stated above. 158 HVGs were calculated in at least 11 of the 22 samples, and these were kept for subsequent dimensional reduction. MultiCCA using 22 dimensions was applied to this batch integrated Seurat object, and the final data set comprised of 10,866 cells. The optimal parameters for resolution set to 1 and perplexity set to 0.75 were selected for FindClusters and RunTSNE, respectively, and this produced 18 communities, which were subsequently re-annotated based on key marker gene expression.

To assign rostrocaudal segment or cell type identity considering the expression pattern of HOX genes (Table S2A) or 105 developmental genes (Table S2B), respectively, all expression values were log transformed after adding a pseudocount of 1. Pearson correlations were performed using pairwise complete observations. Benjamini-Hochberg-corrected p-values for each Pearson correlation were calculated using the corr.test function in the pysch package, and the correlation with the lowest p-value, meeting the specified threshold was used to assign the segment or cell type identity. Multiple identities with the highest correlation were randomly selected for assignment.

### Differential expression, gene set enrichment, and classifier accuracy analyses

To perform differential gene expression analysis between any two populations, the FindMarkers command in Seurat was applied with the bimodal expression likelihood test and the log fold change threshold was set to 0.1. Genes with Bonferroni adjusted P-values less than 0.05 were called significantly changed. Additionally, differentially expressed genes between ALS and control conditions were calculated in DESeq (Anders and Huber, 2010) by summing all scRs UMI counts for each gene across the expression matrix for each sample to simulate bulk RNA-seq expression.

Jaccard indices were calculated by tabulating genes called differentially upregulated or downregulated in each ALS to control of isogenic comparison within each experimental batch and intersecting each set of genes among all experimental batches. The Jaccard index is the ratio of the number of intersecting genes divided by the sum of the union of all genes across the two sets being compared. Gene Ontology (GO) analysis was performed on gene lists using official gene symbols for homo sapiens through the DAVID functional annotation chart. The following categories were tested: OMIM disease, GO Term BP direct, GO Term CC direct, GO Term MF direct, BIOCARTA, KEGG, and REACTOME. Thresholds used were minimum count of 2 and EASE score of 0.1, and GO and pathway sets with Benjamini-Hochberg-corrected P-values of less than 0.5 were called significant and reported. The Vennerable package was used to create Euler and Chow-Ruskey plots. WGCNA and module preservation was performed as previously described in Ho et al., 2016.

Multiset enrichment analysis was performed using SuperExactTest (Wang et al., 2015). Lists of gene sets to be intersected were input, the expected and observed number of overlaps were calculated, and the P-value indicates the likelihood of overlap among all possible comparisons.

To generate a combined expression score for each gene within a population of cells in each sample, we calculated the average UMI counts using all cells within a specified population with a non-zero UMI value. For each gene, the minimum average UMI count among all 22 samples was subtracted from the average UMI in each sample to floor the average expression to zero. From this transformed set of values, the maximum was identified and subsequently used as a divisor for each floored average expression to transform them into a ceiled average expression, with the maximum value being 1. For each gene in each population in each sample, the ceiled average expression was summed with the percent UMI counts for all cells within the specific population, which includes zero UMI values, to generate a combined expression score that equally weights average UMI expression with percent UMI expression. This combined expression score was used to perform statistical test for changes in distribution between all ALS samples and all control and isogenic samples across all experimental batches.

To define MN-specific marker genes for ALS classification, a table of combined expression scores were generated from iPSC-MN scRs data, which contained 1,281 genes. A t-test was performed between all ALS and all control and isogenic samples; 39 genes obtained nominal p-values less than 0.05, and none of these retained this status after Benjamini-Hochberg correction. Therefore, the genes were ranked from lowest to highest nominal p-values, and the top 20 genes were selected to be intersected with the LCM MN transcriptomic data from Rabin et al., 2010 as analyzed in Ho et al., 2016 as well as Krach et al., 2018. Among these, six genes were concordantly changed between ALS and control conditions in all three data sets; the combined expression score was lower in ALS compared to control and isogenic iPSC-MNs, and the gene significance to the sALS component was negative in LCM MNs.

To incorporate the six ALS marker genes into a single prediction metric, principal component analysis (PCA) was applied to samples using the expression values of these six genes, and sample coordinates along the first, second, or a sum of both principal components was used as the prediction metric. For analyzing the scRs samples, the combined expression score for the six genes were used as input. For analyzing bulk RNA-seq data, the log transformed expression values for the six genes were used as input. In some data sets, five of the six genes were used; NDUFAF5 was not annotated in Highley et al., 2014; CARS2 was not annotated in Cox et al., 2010, Kirby et al., 2011, and Kiskinis et al., 2014. For Krach et al., 2018, Highley et al., 2014, and Shi et al., 2018, sample coordinates along PC2 were used as the prediction metric. For Fujimori et al., 2018, the signed values for PC2 coordinates of samples were reversed to place control samples concordant with their placement along PC1. Both PC1 and PC2 coordinates were floored to zero by subtracting the minimum of each PC coordinate, and the sum of the floored PC1 and PC2 coordinates were used as the prediction metric. Coordinates along PC1 were used as the prediction metric for all other data sets. The ROCR package (Sing et al., 2005) was used to plot the Receiver Operator Characteristics and calculate the Area Under the Curve (AUC). The p-value of the Wilcox Rank Sum test was used to determine whether the AUC significantly differs from 0.5, the AUC of an uninformative test.

### DATA AND CODE AVAILABILITY

The scRs data generated in this study are available at the Gene Expression Omnibus under the accession code GSE138121. The scripts written for the analysis performed in this study are compiled into a text document and available https://github.com/ritchieho/2020_scRs_iPSC_ALS.

## Acknowledgements

The authors gratefully acknowledge the following: Tania F. Gendron and Leonard Petrucelli for the polyGP immunoassay data, Victoria Dardov and Jennifer Van Eyk for providing mass spectrometry data for ELAVL3; Kathleen Kurowski, Berhan Mandefro, and Dylan West for assistance with experiments and reagent organization; Soshana Svendsen for critical reading and comments on the manuscript. This work was supported by the following grants: ALS Association (J.R., C.N.S.), California Institute for Regenerative Medicine (RN3-06530, R.H.B.), NIA (K99AG056678, R.H.), NINDS (R01NS069669, R.H.B.), NINDS (U54NS091046, C.N.S.), Target ALS (J.R.).

## Author Contributions

Conceptualization: R.H., K.J.K., K.T., R.H.B., and C.N.S.

Methodology: R.H., M.J.W., K.J.K., and J.G.O.

Software and Formal Analysis: R.H., M.J.W., P.M., K.W., K.J.K., J.G.O., M.K., V.M., and I.K.

Investigation: R.H., M.J.W., P.M., K.W., K.J.K., J.G.O., M.K., V.M., M.G.B., D.O., O.A., S.D.G., and S.H.

Resources: R.H., J.R., K.T., R.H.B., and C.N.S.

Data Curation: R.H., M.J.W., P.M., M.K., V.M., and I.K.

Writing: Original Draft: R.H. and C.N.S.,

Visualization: R.H., M.J.W., K.W., K.J.K., J.G.O., D.O., O.A., S.D.G., and S.H.

Supervision: R.H., K.J.K., J.G.O., L.W., J.R., K.T., R.H.B., and C.N.S.

Funding Acquisition: R.H., J.R., K.T., R.H.B., and C.N.S.

---

**Figure S1.**
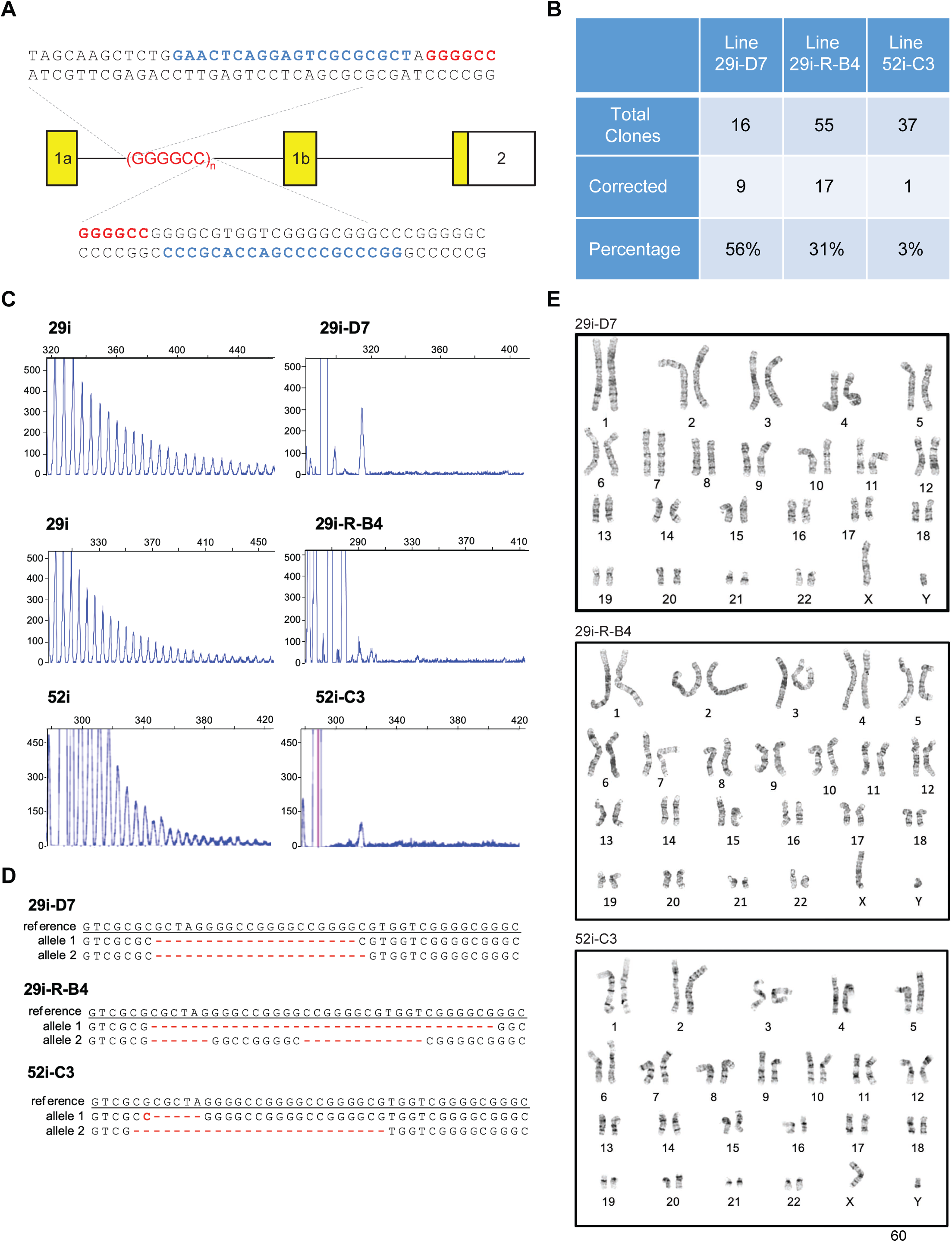
CRISPR-Cas9 removes hexanucleotide repeat expansions (HRE) in two C9orf72 ALS patient iPSC lines. Related to Figure 1. A. Schematic of guide RNA sequences (blue highlights) used to target GGGGCC HRE expansion (red highlight) in the first intron of C9orf72. B. Table summarizing the efficiency of correctly targeted iPSC clones isolated from parental lines 0029 and 0052. 0029-D7 and 0029-RB4 are subclones of 0029, and 0052-C3 is a subclone of 0052. C. Repeat primed PCR assay for HRE in parental and correctly targeted iPSC lines. D. Sanger sequenced C9orf72 alleles from correctly targeted iPSC lines aligned to reference C9orf72 sequence. Red dashes indicate deleted segments, and red nucleotides indicate mismatches. E. Chromosome spreads indicate that the correctly targeted iPSC lines are karyotypically normal.

**Figure S2.**
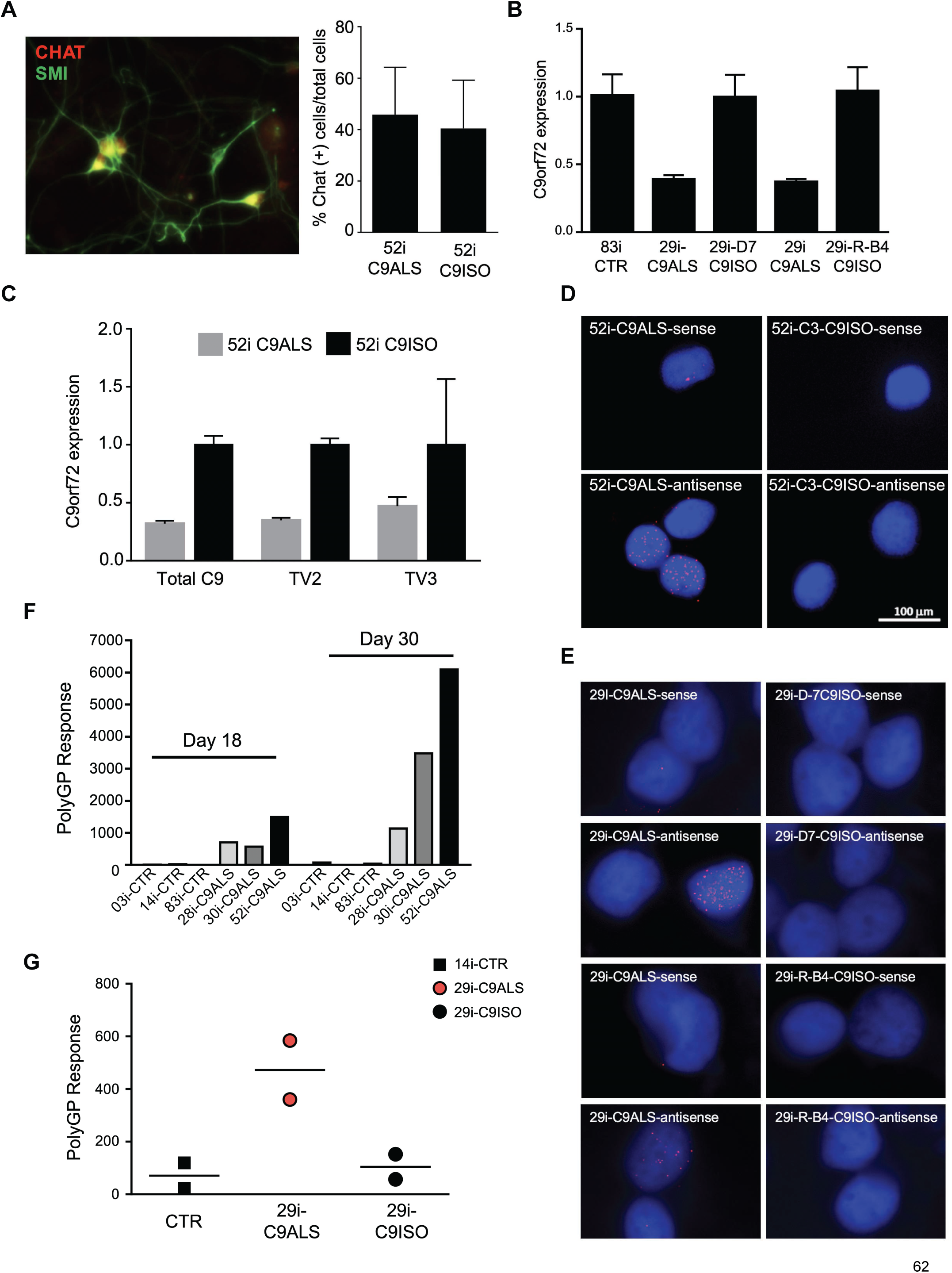
CRISPR-Cas9 corrected iPSC clones differentiate to MNs with altered C9orf72 expression phenotypes. Related to Figure 1. A. Immunostaining MN cultures differentiated from the 0052 corrected iPSC line for the expression of MN markers CHAT and SMI-32 (left). Quantification of percent CHAT-positive cells among total cells in 0052 and isogenic HRE-corrected clone. B. qPCR quantification of total C9orf72 transcript expression in MNs differentiated from control (0083), C9orf72 ALS (0029), and isogenic HRE-corrected clones (0029-D7 and 0029-RB4). C. qPCR quantification of C9orf72 total and transcript variants (TV2 and TV3) expression in MNs differentiated from C9orf72 ALS (0052) and its isogenic HRE-corrected clone. D. Sense and antisense RNA foci are present in MNs differentiated from C9orf72 ALS (0052) but eliminated in its isogenic HRE-corrected clone. E. As in D, for 0029 and its two isogenic HRE-corrected clones. F. ELISA quantification of polyGP dipeptide repeats in MNs differentiated over 18 and 30 days from several control (CTR) and C9orf72 HRE ALS (C9ALS) iPSC lines. G. ELISA quantification of polyGP dipeptide repeats in MNs differentiated over 18 days from a control (0014), C9ALS (0029), and its isogenic HRE-corrected clone, demonstrating reduction of polyGP response as a consequence of HRE removal.

**Figure S3.**
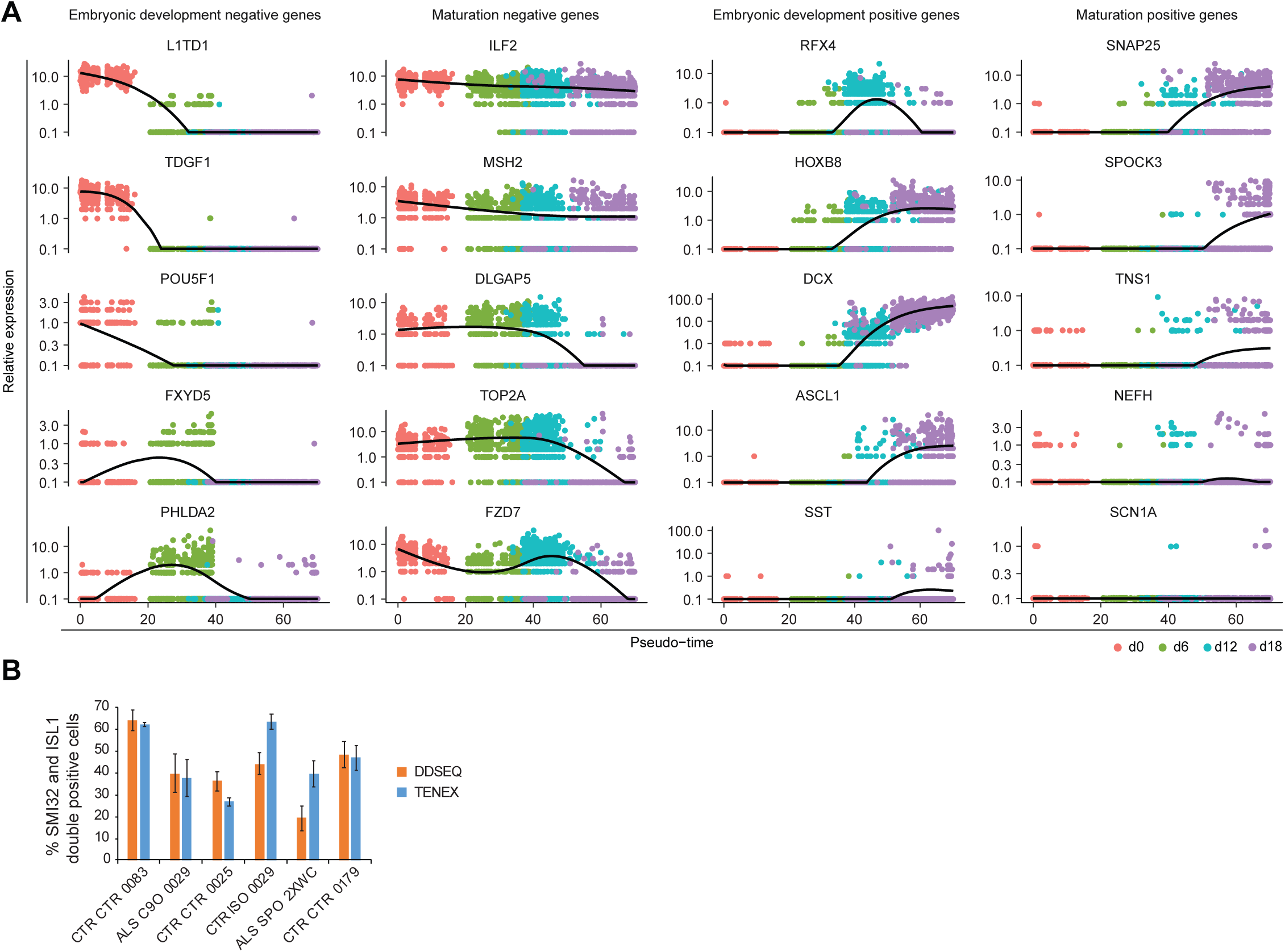
iPSC differentiated spinal MN cultures globally resemble fetal spinal tissue. Related to Figure 1. A. Pseudo-time plots of MN development and maturation markers (Ho et al., 2016) indicate iPSC-MNs recapitulate downregulation of pluripotent stem cell and cell cycle markers and upregulate early spinal MN patterning and progenitor markers with modest expression of maturation markers. B. Quantification of immunostained day 18 cultures from several subject lines (three control, one sporadic ALS, one C9orf72 repeat expansion, and one C9orf72 repeat expansion-corrected isogenic control) for the double positive expression of spinal MN markers ISL1 and SMI-32. Cultures were fixed and stained at day 18, and parallel differentiation cultures were processed on the same day for scRs. Color of bar graphs indicate which differentiation experiments were processed on either the DDSEQ or TENEX scRs technology platform. Three fields of view were quantified and averaged per sample, and error bars indicate standard error.

**Figure S4.**
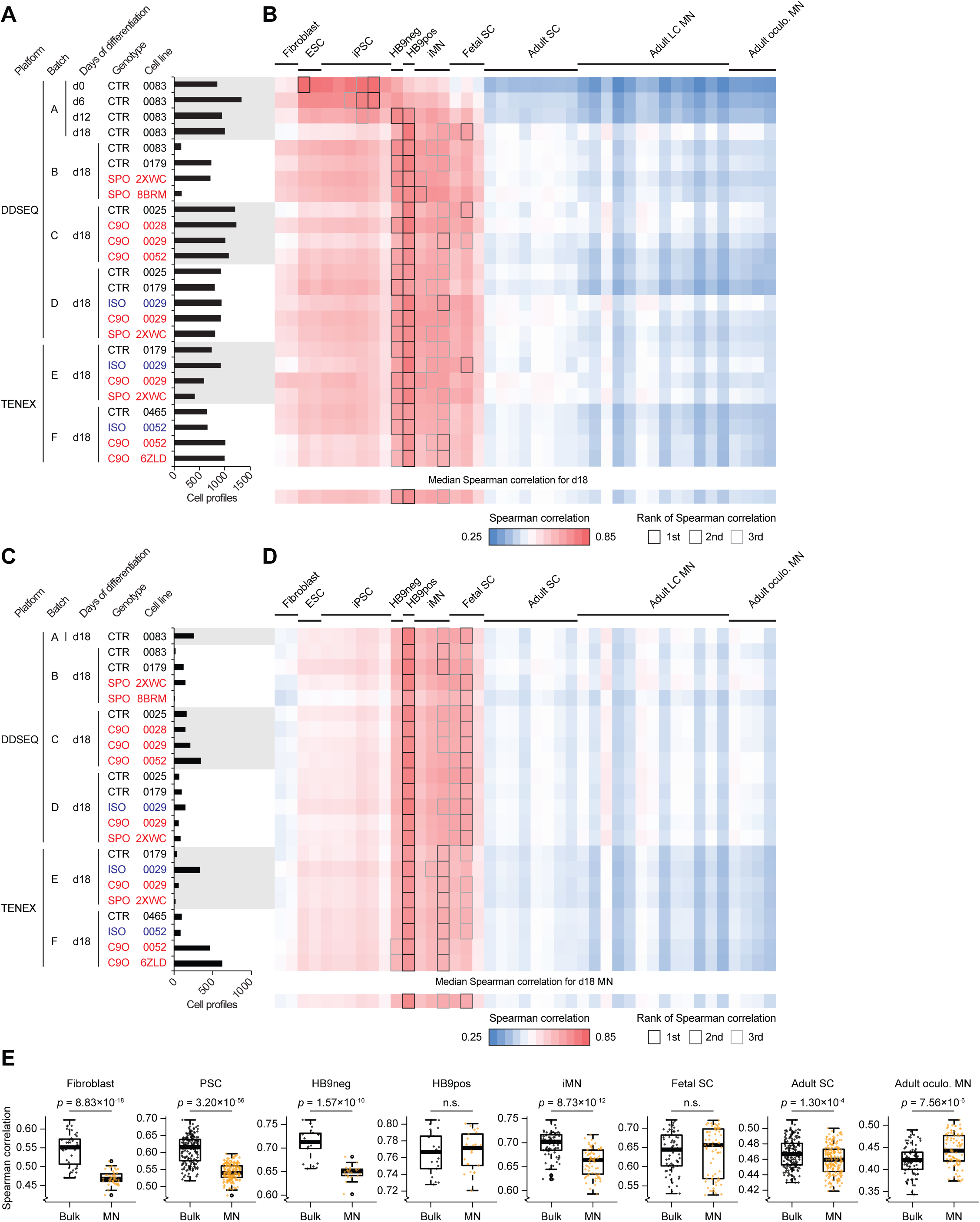
iPSC differentiated spinal MN cultures globally resemble fetal spinal tissue. Related to Figures 1 and 3. A. Histogram displays experimental meta data for all samples profiled using scRs in this study and the number of individual cells that passed quality control filters and subsequently analyzed. This data is also presented in Figure 1. B. scRs UMI counts were summed for each gene across the expression matrix for each sample, and the summed UMI for each gene is meant to simulate bulk RNA-seq expression data for each sample. Simulated bulk gene expression profiles for each sample were correlated to microarray gene expression profiles analyzed in Ho et al., 2016 using Spearman’s method based on the number of matching genes between the two data sets. In comparing the time course experiment from Batch A, there were 9,789 matching genes between the scRs data set and Ho et al., 2016. In comparing the day 18 experiments from Batches B through F, there were 9,989 matching genes between the scRs data set and Ho et al., 2016. The median Spearman correlation was calculated for each column of day 18 samples and displayed along the bottom row. For each row of pairwise correlations, the top three ranked correlations are outlined, indicating which sample from Ho et al., 2016 is most globally similar to each sample analyzed by scRs. ESC: embryonic stem cells, HB9neg and HB9pos: Respectively, GFP-negative and GFP-positive fractions of MN cultures differentiated from ESCs bearing an HB9::GFP reporter, iMN: iPSC-MN cultures, SC: spinal cord, LC MN: laser capture micro-dissected MNs from post-mortem human subjects, Oculo. MN: laser capture micro-dissected oculoMNs from post-mortem human subjects. C. Histogram displays experimental meta data for all day 18 samples profiled using scRs in this study and the number of individual cells classified as MN and subsequently analyzed. D. As in B, for cells from C classified as MNs. E. Boxplots of Spearman correlation values for each sample type in B (Bulk) and D (MN) compared with sampls from Ho et al., 2016. *P*-values were calculated from a paired, two-tailed t-test. n.s.: not significant, *p* > 0.05.

**Figure S5.**
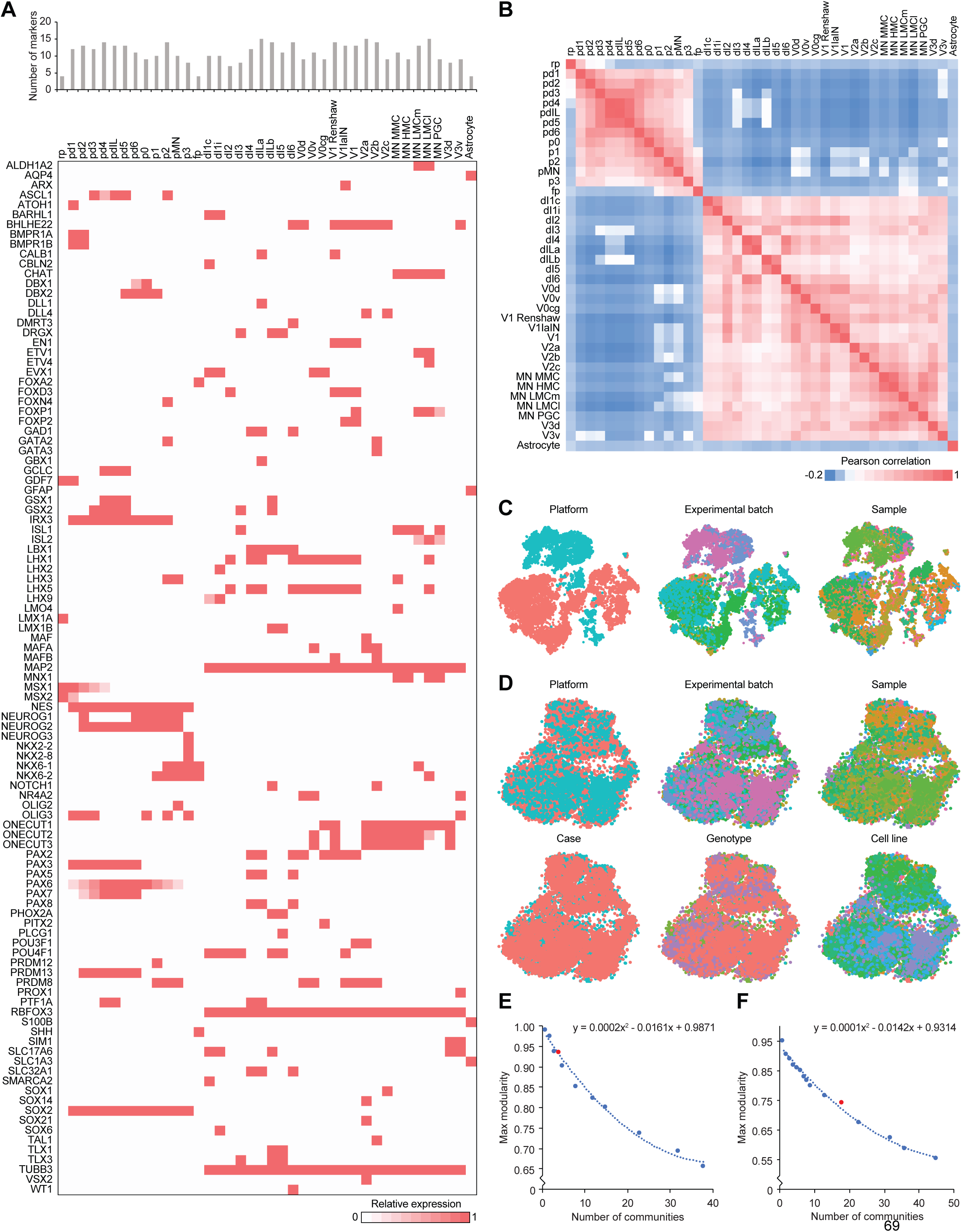
Developmental gene expression profiles distinguish VZ progenitor and MZ postmitotic neuronal identities. Related to Figures 2 and 3. A. Unit expression matrix of 105 genes that developmentally determine VZ progenitor and MZ postmitotic neuronal identities. The number of genes whose expression characterizes each identity is indicated in the bar graph above the heatmap. B. Pairwise Pearson correlation of each identity based on 105 genes in A indicating that the unit expression profile of 105 genes is reasonably sufficient to distinguish each cell type from one another. C. tSNE plot showing global clustering of 17,531 cells from day 18 cultures prior to multiple canonical correlation analysis (Multi-CCA). Each cell is colored by experimental platform (left, red: DDSEQ, blue: TENEX) and experimental batch (middle, n = 5), and sample (right, n = 22). D. tSNE plot showing global clustering of 17,531 cells from day 18 cultures after Multi-CCA. Each cell colored by experimental platform (red: DDSEQ, blue: TENEX), experimental batch (n = 5), sample (n = 22), case (red: ALS, blue: control), genotype (n = 4), and, cell line (n = 10). E. Plot of maximum detected modularity coincident with the number of communities determined by adjusting resolution settings and considering 30 nearest neighbors to construct the shared nearest neighbor graph. The number of communities were determined by testing several resolution settings in the FindClusters command in Seurat. The original Louvain algorithm determined the modularity for each setting, and the maximum modularity observed after 100 iterations was recorded and plotted for each number of communities. A polynomial trendline was calculated (equation is displayed), and the residuals for each setting greater than zero was considered to determine the optimal number of communities. Based on the independently optimized tSNE calculations and visualizations for 17,531 cells, a resolution setting of 0.125 yielding 4 communities was selected to proceed with downstream analysis. F. As in E, except applied to 10,866 classified into clusters 3 and 4 in Figure 3B. Based on the independently optimized tSNE calculations and visualizations, a resolution setting of 1 yielding 18 communities was selected to proceed with downstream analysis.

**Figure S6.**
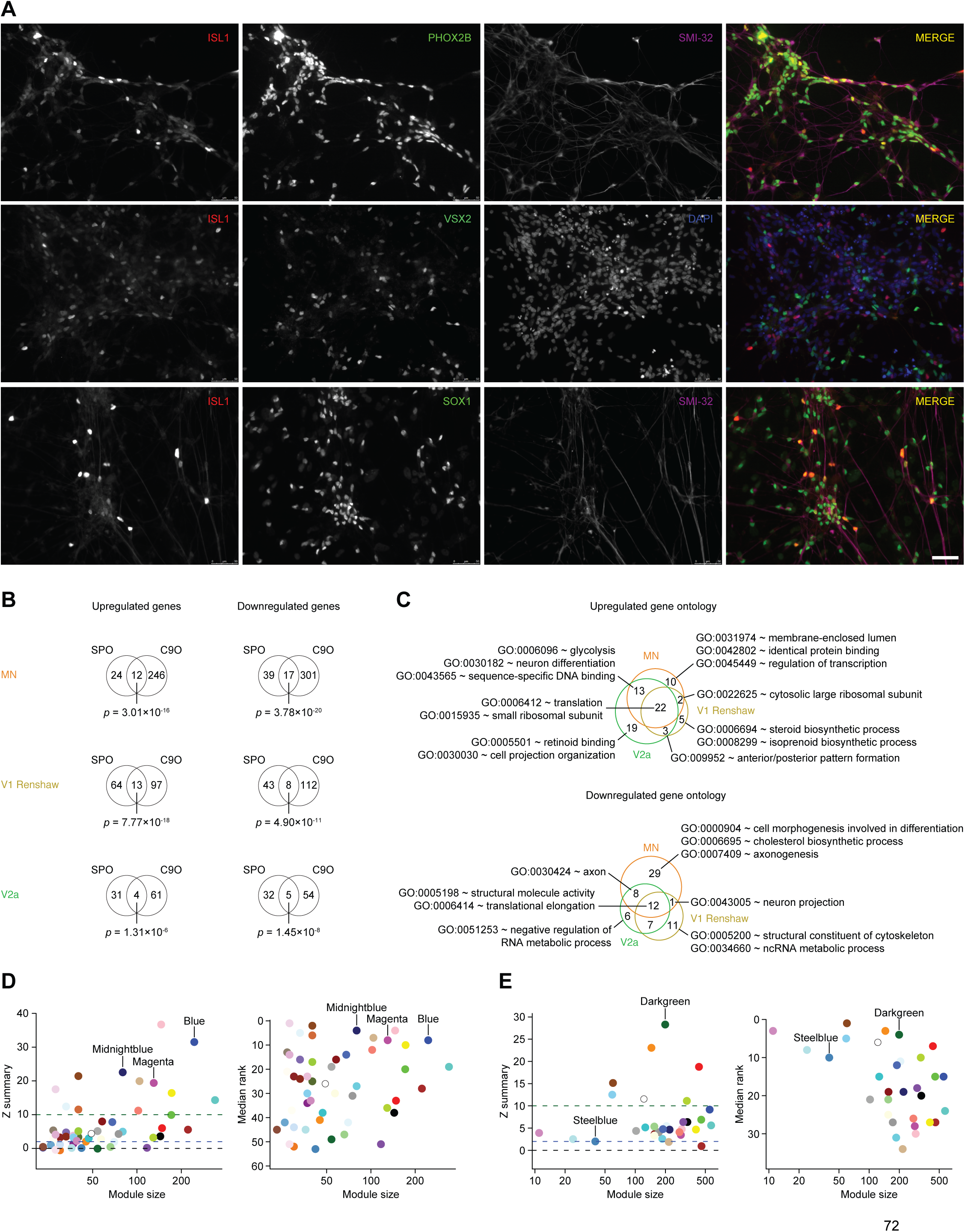
Distinct markers of neuronal subtypes are expressed as proteins in iPSC differentiated spinal MN cultures and enact distinct pathways in ALS conditions. Related to Figures 3, 4, and 5. A. Immunostaining day 18 cultures from one control subject line, 1034 for the expression of spinal MN marker, ISL1, a cranial MN marker, PHOX2B, a V2a interneuron marker, VSX2, and a V2c interneuron marker, SOX1. DAPI and SMI-32 are also used as counter stains. Scale bar, 50 um. B. Venn diagrams of overlapping gene sets between 1) sporadic ALS to CTR and 2) C9orf72 HRE ALS to CTR comparisons within the neuronal subtypes MN, V1 Renshaw, and V2a. Upregulated gene sets were only compared each other, and downregulated gene sets were only compared to each other. P-values of each intersection was calculated by a hypergeometric test. C. Enriched GO terms among genes significantly upregulated or downregulated in ALS compared to control neuronal subtypes are intersect in Venn diagrams. Representative GO terms are displayed from each overlapping or unique set. Full set of GO terms are shown in Table S7B-G. D. Left: measure of how well 52 modules defined in data set A are preserved in the modules defined in data set B. The Z-summary statistic (y-axis) for is plotted against module size (x-axis). Data points reflect module color. Dashed green line indicate threshold at Z = 10, and dashed blue line indicate threshold at Z = 2. For the likelihood of module preservation, Z-summary > 10 indicates strong evidence; 10 > Z-summary > 2 indicates moderate to weak evidence, and Z-summary < 2 indicates no evidence. Right: measure of how well 52 modules defined in data set A are preserved in the modules defined in data set B, as a relative comparison among modules. The median rank statistic (y-axis) is plotted against module size (x-axis). Data points reflect module color. Low median rank values indicate a high preservation. The data points for notable modules featured in Figures 5C and 5D are labeled. E. As in A, except applied to 32 modules defined in data set B and tested for preservation in the modules defined in data set A.

**Figure S7.**
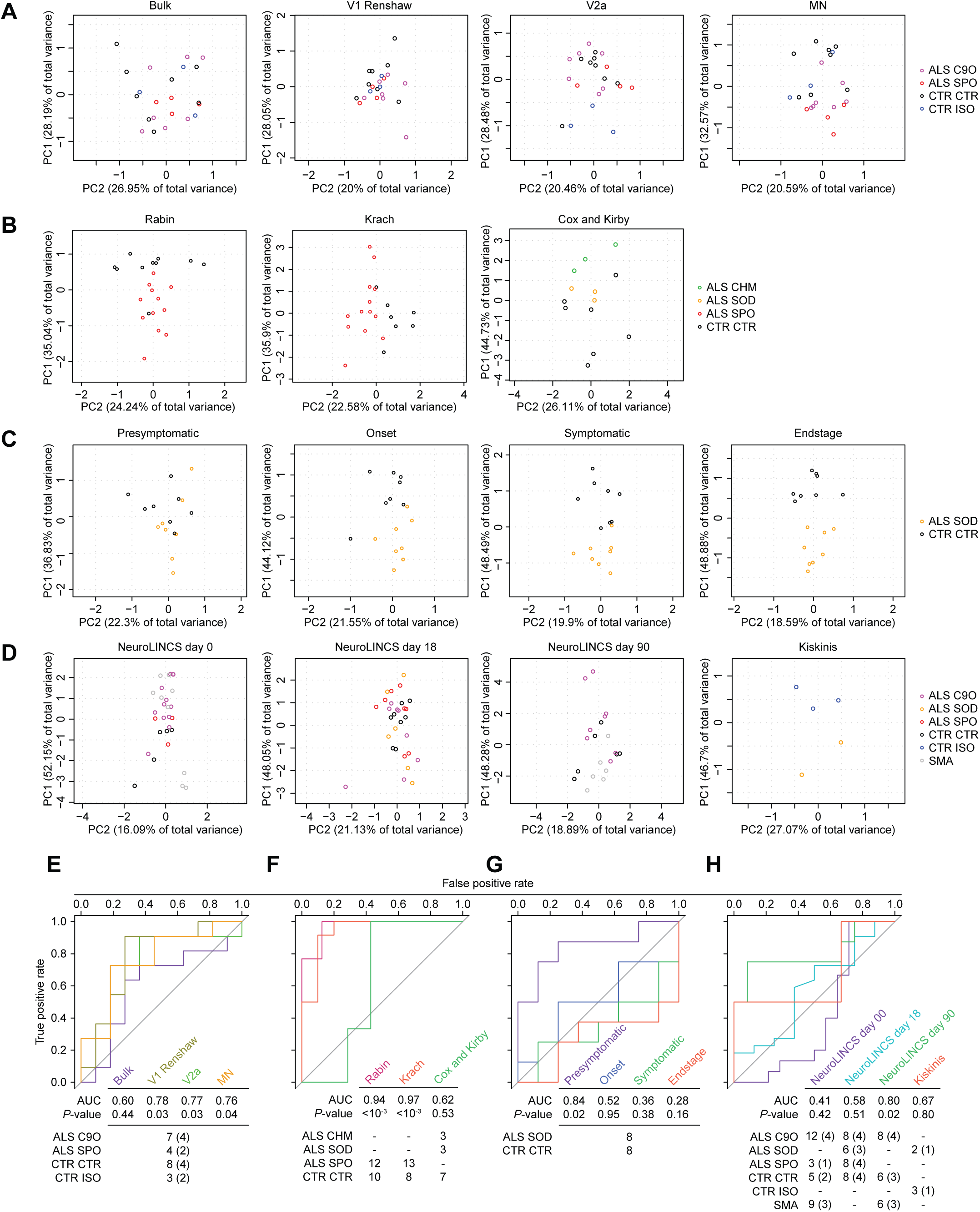
Genes differentially expressed in MN accurately distinguishes ALS from non-ALS conditions throughout the time course of disease. Related to Figure 7. A. PCA on subpopulation data generated from scRs data set based on a combined metric of average and percent expression for the six predictive marker genes: ADCYAP1, CARS2, DNMT3B, ELAVL3, NDUFAF5, and NUAK1. See methods for details on calculation. B. As in A, except on postmortem LC MNs from Cox et al., 2010; Kirby et al., 2011; Krach et al., 2018; and Rabin et al., 2010. For Cox et al., 2010 and Kirby et al., 2011, CARS2 was not annotated and therefore not used in the analysis. C. As in A, except on mouse LC MNs (Nardo et al., 2013). D. As in A, except on bulk RNA-seq data sets generated by the NeuroLINCS Consortium for iPSCs, two MN differentiation protocols at 18 and 90 days of differentiation from iPSCs, and iPSC-MNs profiled by RNA-seq after fluorescent sorting for HB9-RFP-positive MNs (Kiskinis et al., 2014). CARS2 was not annotated and therefore not used in the analysis. E - H. Receiver-operator characteristic (ROC) analysis performed on the data sets described in A-D classifying samples in each data set as ALS or non-ALS. Classifications were based on ELAVL3 mRNA expression. The area under the curve (AUC), which summarizes the overall performance of ELAVL3 expression in correctly classifying samples, is shown beneath each data set. The P-values for each AUC are calculated using the Mann-Whitney U test to determine whether the AUC differs significantly from 0.5 (diagonal grey line), which indicates an uninformative test. The number of samples for each ALS and non-ALS cases used in the analysis, along with their genotypes are indicated. The number of genetically distinct subjects included in the analysis are indicated in parentheses. C9O: familial ALS subjects with pathogenic variants in C9orf72, SPO: sporadic ALS subjects with no known pathogenic variants, CTR: control subjects who do not have ALS, ISO: C9orf72 or SOD1 subject lines in which the pathogenic variant has been genome edited, CHM: familial ALS subjects with pathogenic variants in CHMP2B, SOD: familial ALS subjects with pathogenic variants in SOD1, SMA: spinal muscular atrophy subjects, classified as non-ALS

## Supplemental Table Legends

Table S1. Subject cell line data. Related to Figure 1. Subject clinical and experimental attributes.

Table S2. HOX expression, VZ, MZ, expression, and cell meta data and annotations. Related to Figures 2 and 3. A. Unit expression matrix of HOX genes as they are expressed along the rostrocaudal axis of the developing embryo. B. Unit expression matrix of ventricular and mantle zone developmental marker genes as they are expressed along the dorsoventral and medial lateral axes of the developing embryonic spinal cord. C. Cell meta data and annotations based on rostrocaudal, VZ, and MZ gene expression and global clustering.

Table S3. Differentially expressed genes in MN. Related to Figure 4. Tables of genes called significantly differentially expressed using the bimodal expression likelihood ratio test with log fold-change threshold greater than 0.1 at a Bonferroni-adjusted P-value less than 0.05. Each tab lists significant genes determined from sample to sample comparisons within MN for each experimental batch, separated by upregulated and downregulated categories. ALS to CTR or ISO comparisons were only analyzed between samples within each experimental batch, indicated above each comparison.

Table S4. Differentially expressed genes in V1 Renshaw. Related to Figure 4. Tables of genes called significantly differentially expressed using the bimodal expression likelihood ratio test with log fold-change threshold greater than 0.1 at a Bonferroni-adjusted P-value less than 0.05. Each tab lists significant genes determined from sample to sample comparisons within V1 Renshaw for each experimental batch, separated by upregulated and downregulated categories. ALS to CTR or ISO comparisons were only analyzed between samples within each experimental batch, indicated above each comparison.

Table S5. Differentially expressed genes in V2a. Related to Figure 4. Tables of genes called significantly differentially expressed using the bimodal expression likelihood ratio test with log fold-change threshold greater than 0.1 at a Bonferroni-adjusted P-value less than 0.05. Each tab lists significant genes determined from sample to sample comparisons within V2a for each experimental batch, separated by upregulated and downregulated categories. ALS to CTR or ISO comparisons were only analyzed between samples within each experimental batch, indicated above each comparison.

Table S6. Differentially expressed genes in MNs, V1 Renshaw, and V2a interneurons. Related to Figure 4. Tables of genes called significantly differentially expressed using the bimodal expression likelihood ratio test with log fold-change threshold greater than 0.1 at a Bonferroni-adjusted P-value less than 0.05. Each tab lists significant genes determined from sample to sample comparisons within each of three populations: MN (A and B), V1 Renshaw (C and D), and V2a (E and F) interneurons, respectively separated by upregulated and downregulated categories. ALS to CTR or ISO comparisons were only analyzed between samples within each experimental batch, indicated above each comparison. Genes called significant are highlighted blue. The number of instances each gene was called significant within each category is listed to the right: comparisons, ALS cell lines, CTR or ISO cell lines, and batches. The ratio of each number to total number of comparisons, ALS cell lines, CTR or ISO cell lines, or batches is also displayed. The sum of ratios is calculated to the far right of each table. Genes are displayed based on ranking their ratio sums from highest to lowest. Genes determined to be significant in at least two comparisons were grouped into each set used for subsequent analysis in Figures 4, 5, and 6.

Table S7. Enriched Gene Ontology terms among genes differentially expressed in MNs, V1 Renshaw, and V2a interneurons. Related to Figure 4. A. Differentially expressed genes (DEG) table shows the number of comparisons in which genes were significantly, differentially expressed using the bimodal expression likelihood ratio test with log fold-change threshold greater than 0.1 at a Bonferroni-adjusted P-value less than 0.05. Genes observed to be differentially expressed in two or more comparisons were deemed to be reproducibly dysregulated and thereby kept for subsequent GO analysis. B - G. Tables of Gene Ontology (GO) terms enriched among genes that are reproducibly dysregulated by ALS within each neuronal subtype in the upregulated or downregulated categories. H - M. Tables of Gene Ontology (GO) terms enriched among genes that are reproducibly dysregulated by ALS uniquely within each neuronal subtype in the upregulated or downregulated categories. GO terms are displayed based on ranking Benjamini-Hochberg-corrected P-values from lowest to highest. Benjamini-Hochberg-corrected P-values < 0.05 are highlighted blue. Bonferroni-corrected P-values < 0.01 are highlighted yellow.

